# An integrate-and-fire spiking neural network model simulating artificially induced cortical plasticity

**DOI:** 10.1101/2020.07.23.217265

**Authors:** Larry Shupe, Eberhard E. Fetz

## Abstract

We describe an integrate-and-fire (IF) spiking neural network that incorporates spike-timing dependent plasticity (STDP) and simulates the experimental outcomes of four different conditioning protocols that produce cortical plasticity. The original conditioning experiments were performed in freely moving non-human primates with an autonomous head-fixed bidirectional brain-computer interface. Three protocols involved closed-loop stimulation triggered from (a) spike activity of single cortical neurons, (b) EMG activity from forearm muscles, and (c) cycles of spontaneous cortical beta activity. A fourth protocol involved open-loop delivery of pairs of stimuli at neighboring cortical sites. The IF network that replicates the experimental results consists of 360 units with simulated membrane potentials produced by synaptic inputs and triggering a spike when reaching threshold. The 240 cortical units produce either excitatory or inhibitory post-synaptic potentials in their target units. In addition to the experimentally observed conditioning effects, the model also allows computation of underlying network behavior not originally documented. Furthermore, the model makes predictions about outcomes from protocols not yet investigated, including spike-triggered inhibition, gamma-triggered stimulation and disynaptic conditioning. The success of the simulations suggests that a simple voltage-based IF model incorporating STDP can capture the essential mechanisms mediating targeted plasticity with closed-loop stimulation.

## Introduction

Computational neural network models provide a powerful tool for understanding mechanisms of neural computation and for exploring network behavior in ways that physiological recordings cannot (Fetz, 1993; Fetz and Shupe, 1995; Gerstner and Kistler, 2002; Gerstner et al., 2014). Neural networks consisting of integrate-and-fire (IF) spiking units have proven useful in studying network dynamics produced by spiking neurons [for reviews see (Jolivet et al., 2004; Burkitt, 2006; Gilson et al., 2009a, b, c, d; Ponulak and Kasinski, 2011)]. The IF units typically sum inputs to produce a simulated membrane potential function that triggers a spike when it reaches a threshold. The inputs can simulate post-synaptic potentials with rise time and decay; the units can also incorporate biophysical conductances (Izhikevich, 2003). Spike-timing-dependent plasticity (STDP) rules (Bi and Poo, 2001; Caporale and Dan, 2008; Feldman, 2012; Markram et al., 2012) can be incorporated into IF networks to investigate consequent changes in synaptic connections on network dynamics. For example, large networks of biophysically realistic spiking units with STDP have been shown to form functional interacting groups (Izhikevich et al., 2004; Koene and Hasselmo, 2005; Litwin-Kumar and Doiron, 2014; Bono and Clopath, 2017) and capture global changes induced by conditioning (Song et al., 2013). We here used an IF network with STDP to simulate experimental results from recent physiological conditioning studies performed with a closed-loop brain-computer interface (CL-BCI).

The strength of synaptic connections between motor cortical neurons has been experimentally modified by several different conditioning protocols. In non-human primates (NHP), CL-BCIs have induced plasticity with spike-triggered stimulation of neighboring cortical sites during free behavior (Jackson et al., 2006). After a day of conditioning the output effects on muscles and isometric wrist responses evoked by microstimulation of the recording site (labeled A) included the output effects evoked from the stimulation site (B), consistent with a strengthening of connections from A to B. Changes occurred only for spike-stimulus delays of 50 ms or less, consistent with the time course of STDP. These changes lasted for up to 10 days post-conditioning. A second conditioning protocol triggered cortical stimulation from pulses generated by electromyographic activity of a forearm muscle (Lucas and Fetz, 2013). This produced changes in the output effects evoked by stimulating the cortical site (A) that was associated with the recorded muscle to now include effects evoked from the stimulated site (B). Again, the results were consistent with a strengthening of connections from A to B. A third conditioning protocol used paired sequential stimulation of sites A and B and found changes in the magnitude of potentials at B evoked by stimulation at A (Seeman et al., 2017). These effects were found at some, but not all site pairs, and were again seen for stimulus intervals of 30 ms or less. A fourth conditioning protocol produced cortical plasticity using a CL-BCI to stimulate site A during specific phases of spontaneous beta oscillations at B (Zanos et al., 2018). This produced transient changes in the connection from A to B; the connections increased or decreased, depending on whether stimuli were delivered during the phase in which neurons at B would tend to be depolarized or hyperpolarized, respectively. These changes in connectivity were induced by spontaneous oscillatory episodes with 3 or more cycles and decayed within seconds.

We here investigated whether the neural mechanisms underlying these four conditioning protocols could be captured by an IF neural network model that incorporates STDP. This approach differs from a previous neural network model using populations of Poisson firing units to analytically compute the net effects produced by spike-triggered stimulation (Lajoie et al., 2017) (See Discussion). Our IF model replicated the results of all four conditioning protocols with a single set of network parameters and enabled derivations beyond the original experimental observations, providing a more complete picture of conditioned changes and insights into the effects of relevant network parameters. Furthermore, the model also provided totally novel predictions about the outcomes of possible conditioning experiments that have not yet been performed. Thus, there is a productive symbiotic relation between the model and physiological experiments. Our IF model provides a powerful tool for elucidating the synaptic mechanisms underlying cortical plasticity and discovering new conditioning protocols.

## Methods

### Integrate-and-Fire Network Model

The IF network model consists of 360 integrate-and-fire units. Each unit maintains a potential *V_i_(t)* which represents the sum of the synaptic inputs from units with connections to unit *i* (Fig. 1A). *V_i_(t)* is calculated at discrete time steps of 0.1 ms. When *V_i_(t)* exceeds a threshold θ, the unit “fires” and its spike function *U_i_(t)* is set to 1 for that time step (Eq. 3). Each time a unit fires, its potential is reset to zero on the next time step and an output spike is initiated from that source unit to all its target units.

**Figure 1.**
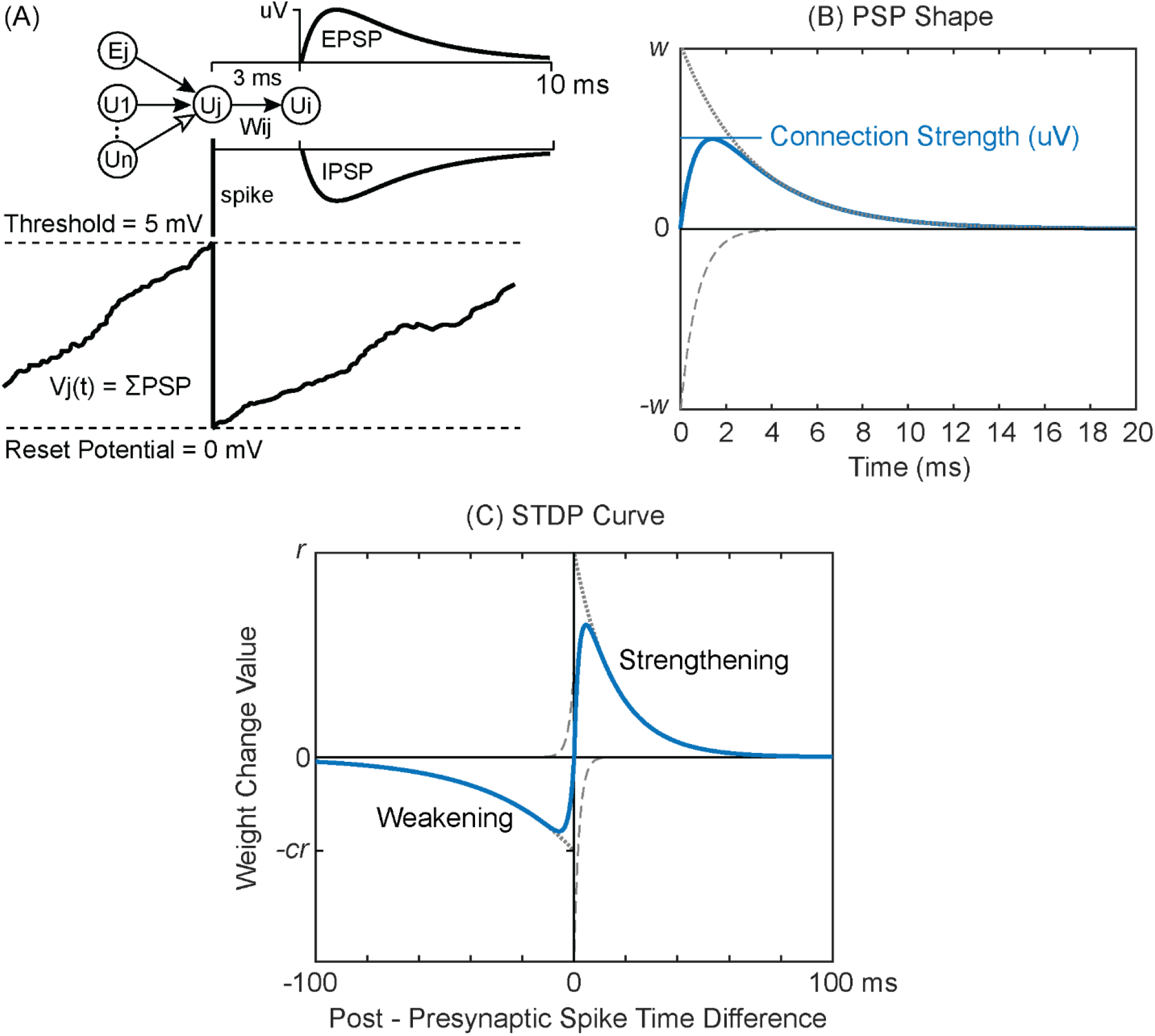
Integrate and fire model. (A) External input (Ej) and spiking input from connected units (U1 to Un) are summed into a target unit’s potential (Vj). When a unit’s potential reaches threshold it is reset to 0 and the unit sends a spike to all its target units. The spike evokes an excitatory or inhibitory post-synaptic potential proportional to synaptic weight *w_ij_*. (B) PSP Shape is calculated as the difference between two exponential functions (dotted and dashed lines). (C) The Spike Timing-Dependent Plasticity Curve shows how much weight *w_ij_* is changed given the difference in spike times between the target unit *i* and the source unit *j*.

The responses of the modeled units to spiking inputs represent the post-synaptic potentials (PSPs) of physiological neurons (Fig. 1B). The form of the PSP is calculated as the difference between two exponential decay functions as follows.

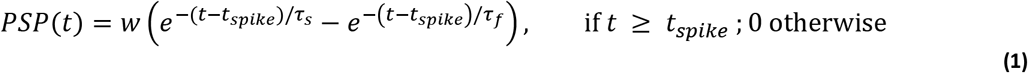

The time constants *τ_s_* (slower decay) and *τ_f_* were chosen to give reasonably shaped PSPs, rising rapidly from zero to a maximum, and then decaying more slowly back towards zero (Fig. 1B). The parameter *w* is the connection *weight –* this parameter is modified by the plasticity calculations. A related parameter, the connection *strength* is measured by the PSP maximum, which represents the synaptic efficacy in voltage change, relatable to distance to threshold. The relation between the weight and strength depends on *τ_s_* and *τ_f_* (Fig. 1B). The calculation of *V_i_(t)* is greatly simplified by using the same value of *τ_s_* and *τ_f_* for all connections. This allows *V_i_(t)* to be represented by two leaky integrators *V^s^_i_(t)* and *V^f^_i_(t)* that decay with these constants (Eq. 4, 5); they both receive the same spiking input and reset with the same spike time function *U_i_(t)*. Here the Euler method is used to estimate the exponential decay functions. We let *A_i_(t)* be the sum of all incoming spike activity to unit *i* at time *t* (Eq. 6), and then calculate the unit potential *V_i_(t)* and spike time function *U_i_(t)* as follows.

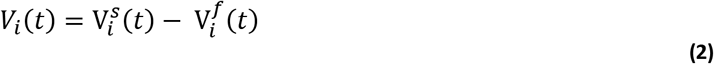

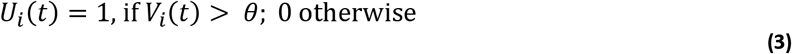

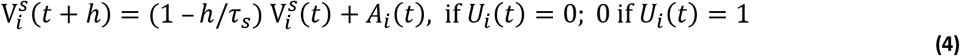

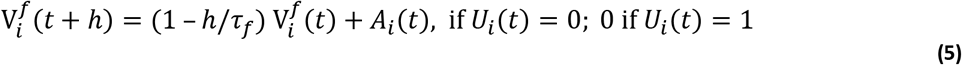

Each connection linking source unit *j* to target unit *i* at time *t* is defined by a weight *w_ij_(t)*. The weight is positive or negative for excitatory or inhibitory units, respectively. Non-existent connections assume a weight of 0. Axonal plus dendritic conduction times are modeled as a global delay parameter *d* for all connections. To provide additional background activity each unit also receives external input *E_i_(t)* the sum of all external spiking activity arriving at unit *i* at time *t* (Eq. 7). *A_i_(t)* is defined as follows:

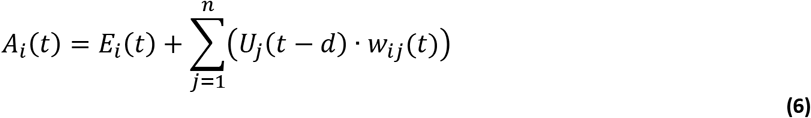

where *n* equals the total number of units. The units are subdivided into three populations called “columns”, representing recorded, stimulated and control sites (see below). To provide background spontaneous activity, each column receives excitatory external inputs *E_i_(t)* that are a combination of correlated inputs *E^c^_i_(t)* and uncorrelated inputs *E^u^_i_(t)*.

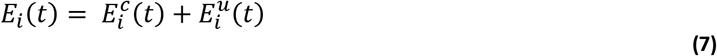

These inputs produce PSPs with the same waveform as other connections in the network, varying only in amplitude. For each column, the correlated events occur with a given probability at each time step; every unit in the column receives the correlated input at a random time within a Gaussian distribution with a standard deviation of 3 ms. Uncorrelated events occur with a separate independent probability at every time step for each unit. The delivery probabilities of correlated and uncorrelated events can be modulated in time to simulate different dynamics of background activity (e.g. oscillations). For our networks *E^c^_i_(t)* and *E^u^_i_(t)* assume a fixed connection weight at times of external spike delivery and are 0 otherwise.

### Plasticity Rule

When spike-timing dependent plasticity is active, connection weights are modified based on the relative firing times of the target and source units. The STDP Curve (Fig. 1C) shows how much weight *w_ij_* changes as a function of the difference in spike times between target unit *i* and source unit *j*. The curve has two components: a strengthening portion for positive differences *Δt* between time of post-synaptic spike and arrival of presynaptic spike that will strengthen the weight, and a weakening portion for negative time differences that will weaken the weight. Each half of the STDP function was approximated with exponential decay functions similar to the PSP function except with longer time constants (*a_s_*, *a_f_*, *b_s_*, *b_f_*) and with amplitudes controlled by the training factor *r* and a weakening factor *c*.

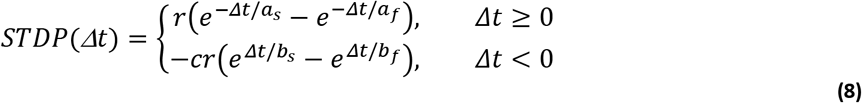

The weakening side of the STDP curve has a lower amplitude but decays more slowly than the strengthening side, and has larger overall area (Caporale and Dan, 2008). The same STDP function was used for excitatory and inhibitory synaptic connections, for which there is empirical support (Haas et al., 2006). For further discussion of this and alternate STDP functions for inhibitory synapses, see Discussion.

Choosing the right amount of weakening vs strengthening is important to the evolution of the network connections. If the weakening factor is too small, the greater area of the strengthening side of the STDP curve will eventually cause all weights to grow to a maximum limit. If the weakening factor is too high, the greater area of the weakening side will push all weights strongly toward zero. We chose a weakening factor of c = 0.55, so that the network sustained low weight values in the absence of any conditioning stimuli but showed noticeable increases in some weights when conditioning was applied.

For a single weight *w_ij_* and time difference between spikes of unit *i* and arriving spikes of unit *j*, the change in *w_ij_* is equal to the value taken from the STDP function in Figure 1C. To facilitate computation of weight changes, strengthening and weakening potentials are maintained for each unit similar to the way unit potential *V_i_(t)* sums PSPs (except with no threshold crossing reset). *S_j_(t)* sums the strengthening side of the STDP curve for each source unit *j* using time constants *a_s_* and *a_f_* with input activity *U_j_(t–d)*. *T_i_(t)* sums the weakening side for each target unit *i* using time constants *b_s_* and *b_f_* with input activity *U_i_(t)*. Thus, when a spike occurs, weight changes can be calculated with respect to previous spikes by taking signed and scaled values of *S_j_(t)* and *T_i_(t)* (Eq. 15).

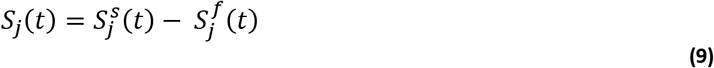

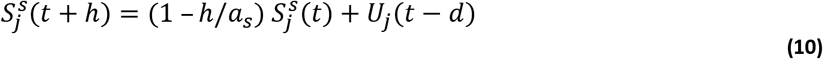

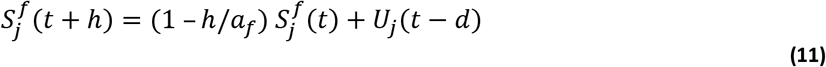

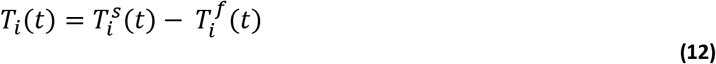

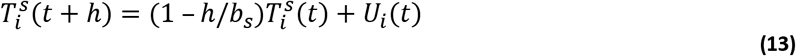

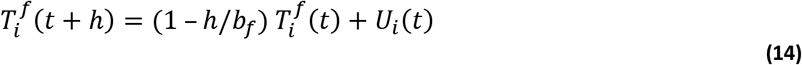

The change to *w_ij_* at a given time step is:

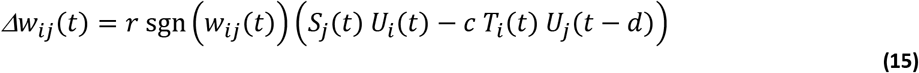

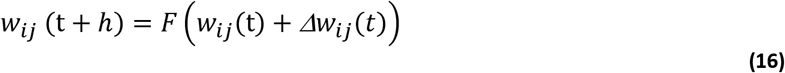

The function *F(w)* clips excitatory weights to the range *w_min_* ≤ *w* ≤ *w_max_*, and inhibitory weights to the range −*w_max_* ≤ *w* ≤ −*w_min_*. This function can be modified to squash weight changes as the weight approaches min and max limits, but for the simulations here no graded squashing was performed. The standard parameter values used in simulations are shown in Table 1.

**Table 1.** Parameter values.

|  |  |
| --- | --- |
| Number of excitatory units in each column | 40 |
| Number of inhibitory units in each column | 40 |
| Unit firing threshold | 5 mV |
| Conduction delay between cortical units | 3 ms |
| Conduction delay to motor outputs | 10 ms |
| External Input PSP strength | 350 μV |
| Unit potential time constants $\tau_s, \tau_f$ | 3.2 ms, 0.8 ms |
| External input rate | 1800 spikes per second |
| External input correlated spike percentage | 30% |
| External input correlated spike standard deviation | 3 ms |
| Training factor $r$ | 100 |
| Weakening factor $c$ | 0.55 |
| STDP strengthening curve time constants $a_s, a_f$ | 15.4 ms, 2 ms |
| STDP weakening curve time constants $b_s, b_f$ | 33.3 ms, 2 ms |
| Conditioning stimulus intensity | 2 mV |
| Maximum connection strength (used to calculate $w_{max}$) | 500 μV |
| Minimum weight $w_{min}$ | 1 |
| Weight change squashing (weight dependence) | 0 (none) |
| Probability of excitatory cortical connections | 1/6 |
| Probability of inhibitory connections | 1/3 |
| Probability of corticomotoneuronal connections | 1/3 |
| Number of motoneurons | 40 |
| Motoneuron thresholds | 5 to 6 mV |
| Motor unit potential size | 0.5 to 1.5 mV |
| Available simulation time step sizes | 0.1 (default), 0.05, 0.025, 0.02, 0.01 ms |

### Network Topology

The network contains 240 cortical units grouped into three “columns”, A, B and C, as shown in Figure 2A. Each column has 40 excitatory units (e) that project only positive weights, and 40 inhibitory units (i) that project only negative weights. Excitatory units connect sparsely to all other units in the network and inhibitory units connect less sparsely to all units within the same column. The probability of each possible connection is listed in Table 1. Unit self-connections are not allowed. Each column also has a simulated local field potential (LFP) which is the sum of all post-synaptic potentials occurring within a column.

**Figure 2.**
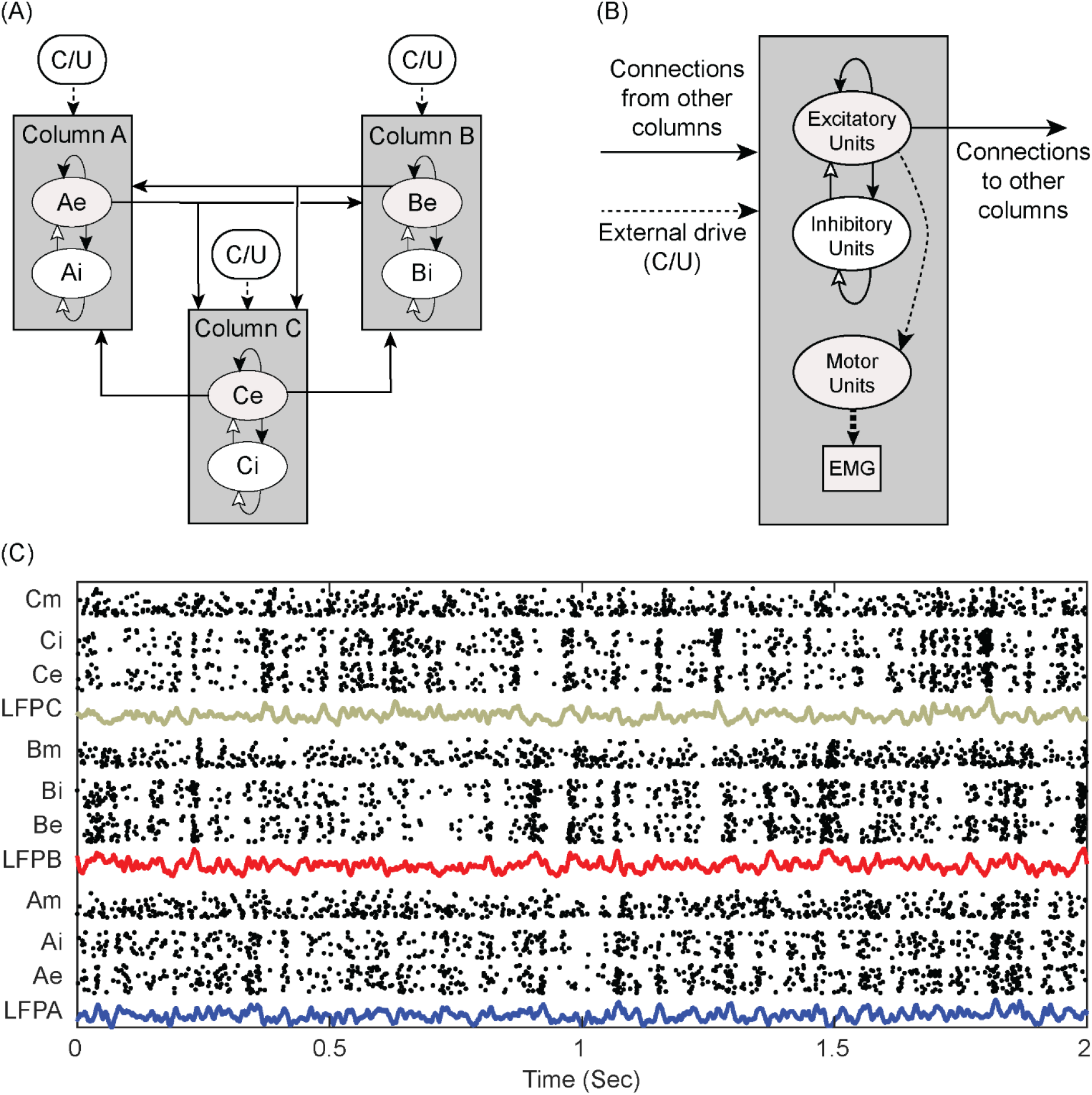
Network Connectivity. (A) Cortical network configuration. Units are grouped into 3 columns A, B, and C. Each column contains 40 excitatory units (e.g. Ae1 … Ae40) and 40 local inhibitory units (Ai) all sparsely interconnected. Only excitatory units project to other columns. Columns also receive external input, consisting of correlated and uncorrelated (C/U) exponentially distributed spikes. (B) Column configuration. Associated with each cortical column is a pool of motor units that produce muscle potentials (EMG). The dashed lines designate unmodifiable connections. (C) Example spiking activity of excitatory (e), inhibitory (i) and motor units (m) associated with each column. LFP is the sum of within-column post-synaptic potentials in e and i units.

To provide background spontaneous activity each column receives an external excitatory bias input generated separately for each column. Some bias inputs provide correlated spikes to each unit in a column, and others provide independent uncorrelated spikes to all units (C and U in Figure 2A), for a combined mean bias rate of 1800 spikes/sec to each unit. The ratio of the number of correlated to uncorrelated spikes each unit receives can be modified to control the degree of synchronization within a column. The mean bias rate can also be modulated over time, for example to generate oscillatory activity or simulate behavioral activation.

To simulate experiments that involved recording muscle activity, the network includes pools of motor units driven by the cortical columns (Fig. 2B). Associated with each column is a pool of 40 motoneurons that receive inputs from a third of the excitatory units of that column, as well as from uncorrelated external drive to simulate all additional inputs. Consistent with the size principle (Henneman et al., 1965), the pool included a range of small to large motoneurons, with increasing thresholds from 5 to 6 mV and increasing muscle unit potentials from 0.5 to 1.5 mV. Multiunit electromyographic (EMG) activity was simulated by summing all the muscle unit potentials in the same manner as for unit PSPs and then filtering the result with a 100 to 2500 Hz bandpass filter.

Figure 2C shows the simulated spiking activity of all units under steady-state conditions. Also plotted are the corresponding LFPs for each column. The coordinated bursts of activity are produced by 30% correlated external bias drive.

### Conditioning Stimulation

The effect of electrical stimulation on a stimulated unit is modeled by a large and immediate deflection of its potential towards threshold, computed by adding a large stimulus pulse to the unit’s *V^s^_i_(t),* proportional to stimulus intensity. The standard conditioning stimulus produces a step in *V_i_*(t) of amplitude 2 mV. The standard testing stimulus for evoking potentials produces a step in *V_i_*(t) of amplitude 3 mV. Normally a conditioning stimulus is applied to all units in a column simultaneously, causing many of them to fire. This burst of spikes will evoke a measurable response in the LFP of other columns, called the Evoked Potential (EP). The EP is a measure of the net synaptic strengths from the stimulated column to the recorded column. The size of each EP is calculated as the difference between the amplitude of the LFP peak after the stimulus compared to pre-stimulation baseline LFP. The average change in the EP amplitude produced by conditioning is quantified as the percent increase of the average EP amplitude after conditioning compared to before conditioning (Eq. 17).

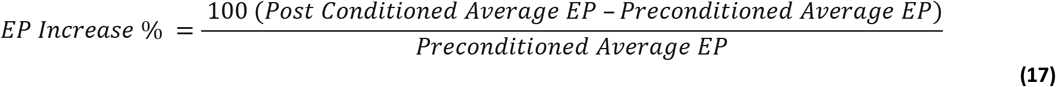

### Implementation

The network has many modifiable parameters that can affect the outcome of a simulation (Table 1). Temporal parameters are generally scaled in milliseconds. The network runs in time steps of 0.1 ms to strike a balance between accuracy and computation time. Connection strengths, stimulation amplitudes, and PSP-related parameters are scaled in microvolts to relate such values to physiological variables. Connection strengths are initially small values (20 to 60 percent of maximum) taken randomly from a uniform distribution.

Most networks were run in 10-second time blocks for 2000 seconds of simulated time, divided into four periods of 500 seconds each (Table 2). During the preconditioning period STDP is on and the network settles into an unconditioned steady state. During the two testing stages before and after conditioning the STDP calculations are turned off, allowing graphs and tests to be compiled while the network runs with static weights. The conditioning protocol is run during the conditioning period with STDP on.

**Table 2.** Sequential training periods.

| Period | Preconditioning | Preconditioning<br>Testing | Conditioning | Post-conditioning<br>Testing |
| --- | --- | --- | --- | --- |
| Plasticity | On | Off | On | Off |
| Conditioning | Off | Off | On | Off |

### Tetanic Stimulation

To control for the effects of stimulation alone, tetanic stimulation was performed with a 10 Hz Poisson spike train with an imposed 10 ms refractory period. This can have a strengthening effect on connections from the stimulated column to the other two columns, and a weakening effect in the reciprocal directions. These effects were minimized by selecting a weakening factor large enough to yield preconditioned networks with small weights that would readily show the effects of conditioned strengthening (Fig. 8A). There is substantial variation in the sizes of conditioned EPs (Fig. 8B) due to differences in initial conditions, the randomness of the external input, and the magnitude of the training factor parameter. However, averaging over a set of simulations using 10 different initial conditions yields a reliable measure of the effects of tetanic stimulation (Fig. 8C). These tetanic effects can support or oppose other conditioning. For example, any protocol to strengthen A→B connections that employs stimulation on Column B must overcome the tendency for tetanic stimulation on B to weaken those connections. On the other hand, protocols that stimulate Column A (like cycle-triggered stimulation) should be evaluated relative to simple tetanic stimulation of Column A.

### Code Accessibility

The Matlab [v. 2019a] code for the IF network model is available online at https://github.com/lshupe/spikenet. Online material includes documentation and instructions on running the model under Windows 10, as well as further explanation of effects of network parameters. The code includes options not exercised in these simulations, such as including a squashing function for graded weight changes and running with finer temporal resolution (which did not significantly alter the main results).

## Results

### Spike-Triggered Stimulation

To simulate conditioning with spike-triggered stimulation (Jackson et al., 2006; Rebesco et al., 2010), each time the first excitatory unit (Ae1) in Column A fired, a conditioning stimulus was applied to all units in Column B at a given delay (Fig. 3A). Conditioning effects were observed as increased connection strengths between Column A and Column B. Figure 3C shows color-coded strengths of all connections after spike-triggered stimulation with a spike-stimulus delay of 10 ms, showing strengthened connections from Ae to B units compared to connections between other columns. Connectivity between Columns A and B was also documented by the average evoked potential (EP) (Fig. 3D insets) in the LFP of Column B produced by a test stimulus to A (Fig. 3A).

**Figure 3.**
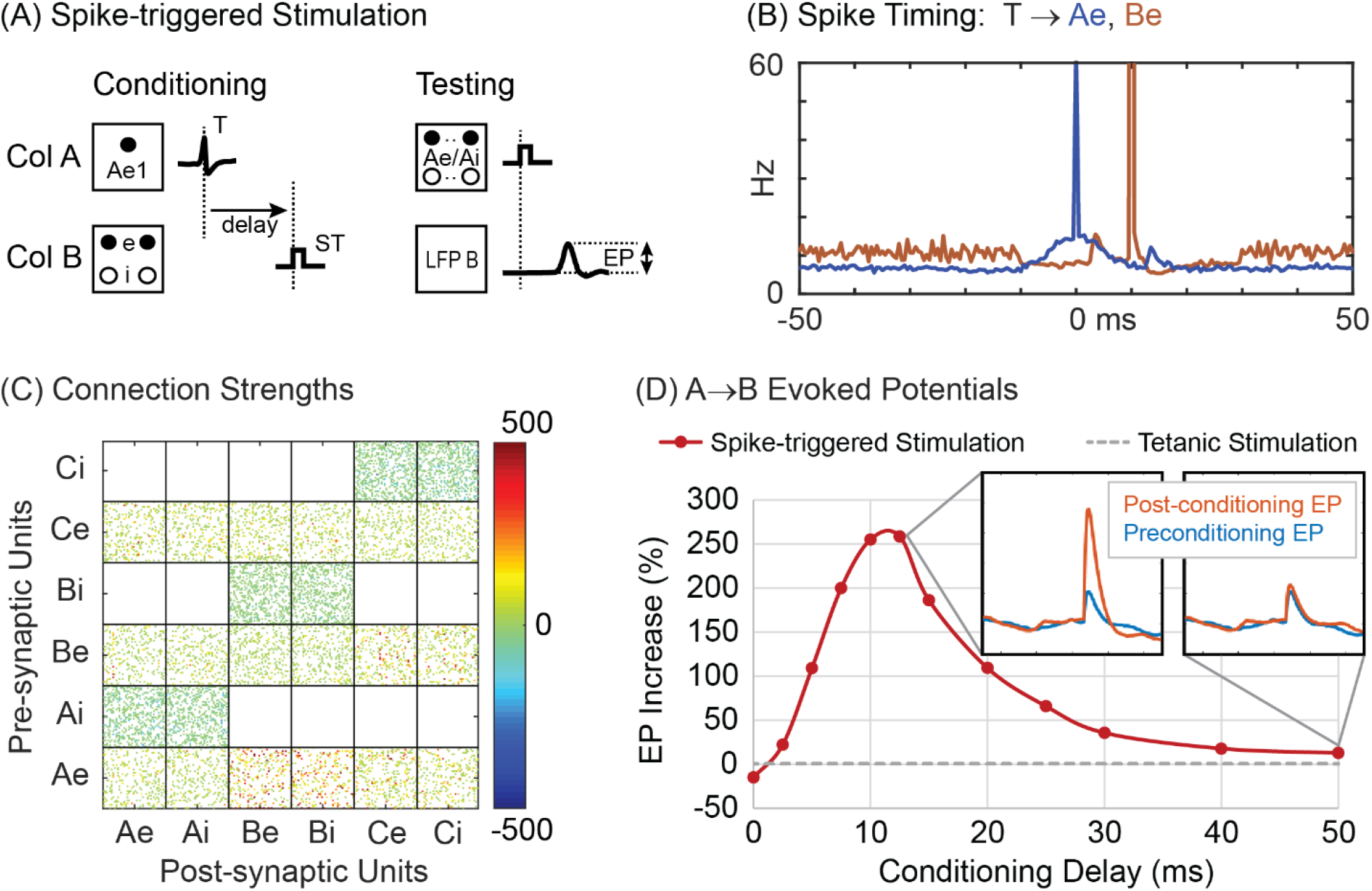
Spike-triggered stimulation. (A) A Conditioning stimulus (ST) is applied to all units in Column B at a delay from each trigger spike (T) detected on the first excitatory unit (Ae1) of Column A. Testing shows stimulus applied to all Column A units to evoke a potential (EP) in LFP of Column B. (B) Histogram of excitatory A and B unit firings aligned with spikes on Ae1 (T) during conditioning with a 10 ms delayed stimulus. (C) Final connection strengths (calibrated in μV) between units after conditioning. (D) Average percent increase in the EP as a function of the delay between trigger spike and stimulation. Insets show average EPs before and after conditioning for two delays. Grey dashed line shows the EP response in Column B after tetanic stimulation of Column B.

After conditioning for 500 seconds of simulation time, the conditioning unit (Ae1) showed near maximum connection strengths to its target units in Column B. Other units in A also show increased connections to B depending on the percent of correlated bias inputs to A. Networks with 30% correlated bias inputs show a moderate conditioned response in other Column A→B connections. Networks with 20% correlated bias inputs show little conditioned response except for connections from unit Ae1. Figure 3B shows the average firing rates of the Ae and Be populations relative to the triggering Ae1 spike times (T). The sharp peak at 0 ms represents the trigger spikes and the broad peri-trigger peak reflects the synchrony between Ae units imposed by 30% external correlated bias. Figure 3B also shows the times of spikes in Column B (red); the large peak at 10 ms reflects the occurrence of stimulus pulses in B.

Figure 3D shows the size of A→B EPs after spike-triggered stimulation at various spike-stimulus delays. The peak in the curve reflects the effect of the STDP rule. At long delays, the spike-triggered conditioning effect approaches the small effect of tetanic stimulation applied to Column B (dashed line). At “zero” delay the size of the EP is decreased (Fig. 3D). In this case the spike-stimulus delay is shorter than the conduction delay and connections from A to B become weaker. This decrease in synaptic strength is consistent with the STDP rule and with experimental results obtained for corticospinal connections (Nishimura et al., 2013).

The original experiments of Jackson et al (Jackson et al., 2006) could not document the conditioning effects by recoding short-latency EPs because of stimulus artifacts, so instead measured conditioning effects indirectly by using cortical microstimulation to evoke EMG responses. To simulate these experimental observations, the motor unit pools were activated by trains of cortical stimuli delivered at separate times to each column (Fig. 4). This simulation shows that after conditioning, stimulation of Column A now also evoked responses in the muscle of Column B, mediated by the strengthened A→B connections and as reported by Jackson et al (their figure 2).

**Figure 4.**
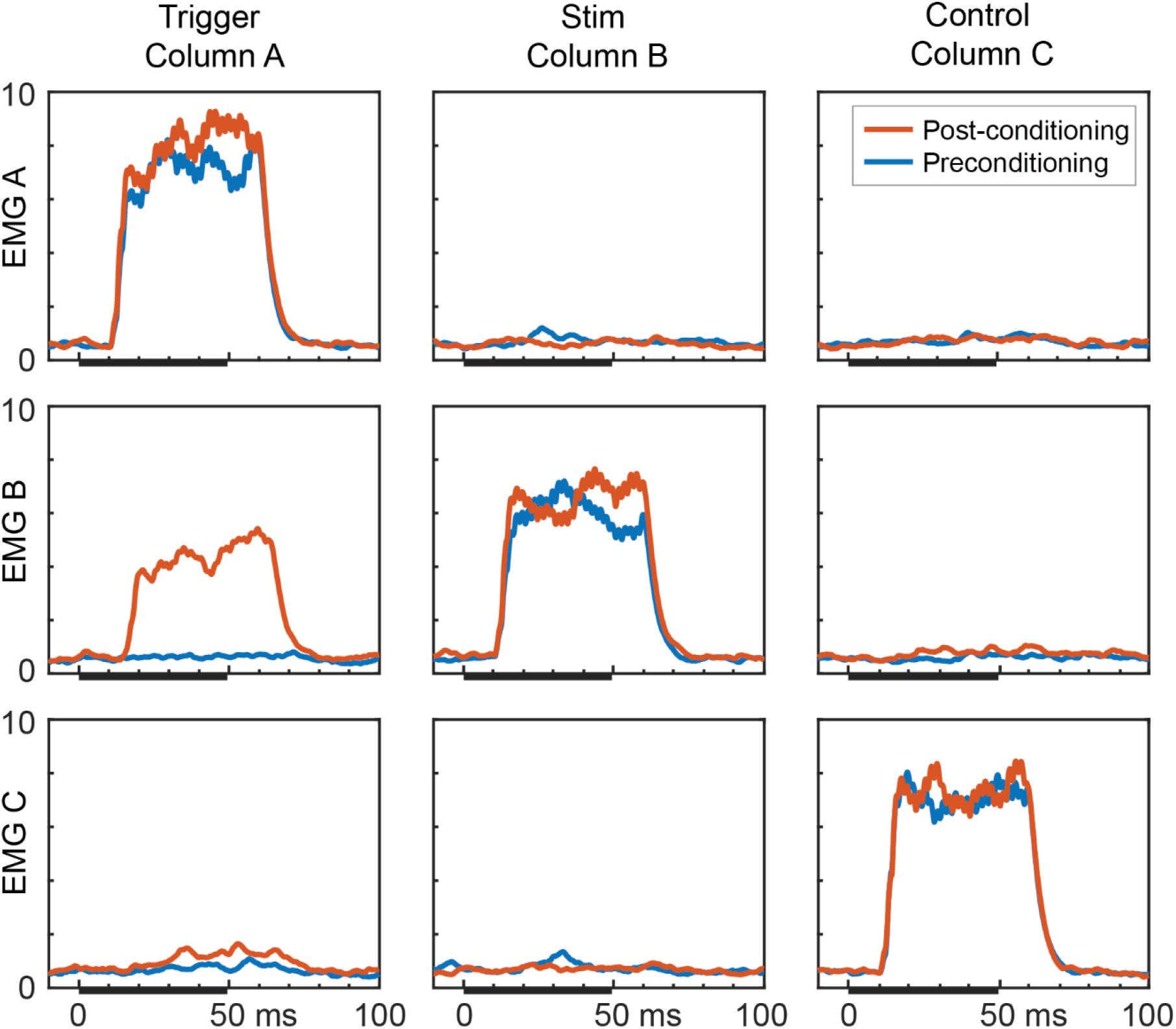
Simulation of output effects on muscles before and after spike-triggered stimulation. Averages of rectified EMG responses evoked by repetitive stimulation of Column A (containing conditioning trigger unit), Column B (stimulus column), and Column C (control) for preconditioning (blue) and post-conditioning (orange) periods. [cf. figure 2 in (Jackson et al., 2006)] For this simulation, inter-column connection probability was doubled to 1/3 and the correlated bias drive was set to 50%. The test stimulus train was 1 mV, 25 pulses, 500 Hz delivered during intervals marked by black bars (the response delay is due to the 10 ms corticospinal conduction time). Ordinate scale is in millivolts.

### EMG-Triggered Stimulation

Cortical conditioning effects could also be produced in NHPs by triggering cortical stimulation from muscle activity (Lucas and Fetz, 2013). To simulate these experimental results the trigger pulses were obtained from threshold crossings of multiunit EMG activity of Muscle A (Fig. 5A). The threshold was chosen such that triggered stimuli were delivered to Column B at a rate comparable to that used for other conditioning methods (approximately 3000 conditioning stimulations over a 500 second conditioning period). This protocol strengthened the synaptic connections from A to B, as shown by the connectivity matrix (Fig. 5C) and by the A→B evoked potentials (Fig. 5D). The conditioning effect is explained by the computed histogram of spikes in the Ae units aligned with the EMG threshold detection (Fig. 5B). This shows a peak in firing of A units (blue curve) that generated the coincident input to the motor units and that preceded the triggered responses in B (red peak). The delay between these peaks represents the 10 ms cortico-motor-unit conduction time plus a minor effect of EMG rise to trigger threshold.

**Figure 5.**
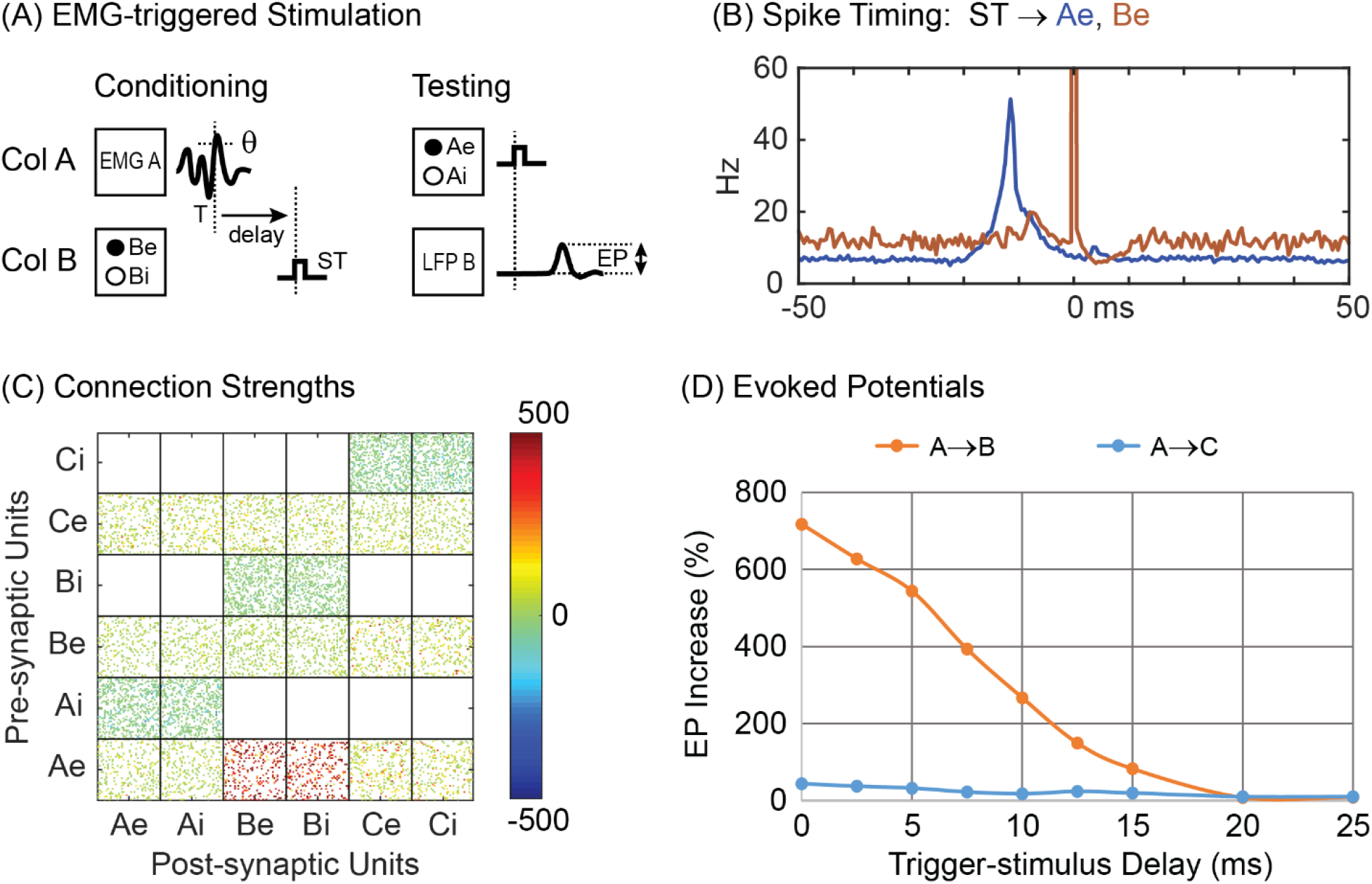
EMG-triggered Stimulation. (A) A conditioning stimulus is applied to all units in Column B at a delay after threshold crossing (θ) in EMG of A motor units. (B) Firing rate histograms of Ae and Be units during conditioning aligned with EMG A threshold detection and stimulation (0 ms). (C) Connection strengths between pre and post synaptic units after conditioning. (D) The average percent increase in the EPs evoked in Column B (orange) and C (blue) from stimulating A as function of delay between EMG threshold crossing and delivered stimulation.

### Paired Pulse Stimulation

Paired pulse conditioning (Rebesco and Miller, 2010; Seeman et al., 2017) was simulated by stimulating Column A followed by stimulation of Column B at a fixed delay (Fig. 6A). The resultant connection strengths are shown in Figure 6C for a stimulus delay of 10 ms. Conditioning effects were also documented by delivering test stimuli to Column A and measuring the average EPs in Columns B and C. Figure 6D plots the change in amplitudes of the A→B EPs and A→C EPs as a function of interstimulus delay. Consistent with the bidirectional STDP function, there is a decrease in the size of the EPs for negative conditioning delays (i.e. B stimulated before A), but this is shallower than the peak for positive delays because of the choice for the weakening parameter, which tended to keep synaptic weights small. The histograms of unit firings shows the peak in A spikes produced by the stimulus in A (blue trace) and two peaks in B firing (red trace): the first peak is due to a synaptically relayed response to the A burst and the second is due to the delayed stimulus of B.

**Figure 6.**
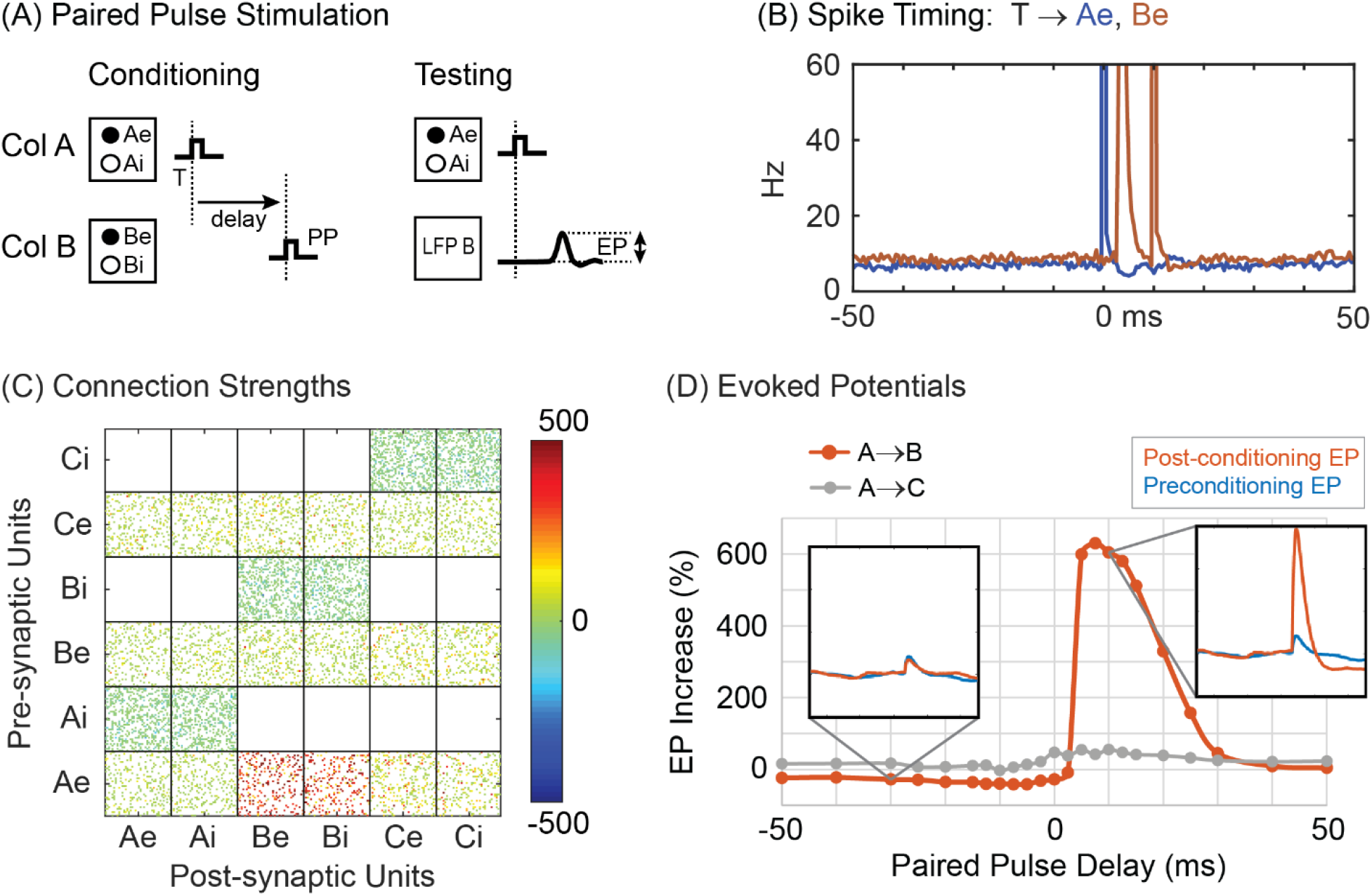
Paired pulse stimulation. (A) Pairs of pulses delivered sequentially to A and B with fixed delay. (B) Firing rate histograms of Ae and Be units during conditioning aligned with times when the first of the paired stimuli occurred (10 ms conditioning delay). (C) Connection strengths after conditioning. (D) Average percent increase in the evoked potential between the conditioned and unconditioned effect as a function of the interstimulus delay. Insets show the shape of the average EP at conditioning delays of −30 ms and 10 ms. Grey trace plots same for EPs in Column C.

For the same stimulus amplitude, paired pulse conditioning tends to be stronger than spike-triggered conditioning (see Testable outcomes and Figure 10) and does not require correlated bias inputs, since the stimulation pulses themselves evoke strong correlated activity between A and B. The conditioning effect is seen with far fewer stimuli than used for spike-triggered stimulation, for the same stimulus intensity. Our 500-second conditioning time yielded only 700 paired pulse conditioning stimuli, compared to over 3000 stimuli when using the spike-triggered stimulation protocol.

In these simulations conditioning effects were obtained by delivering pairs of single pulses. However, in the physiological experiments it was necessary to use a triplet of pulses to obtain effects (Rebesco and Miller, 2010; Seeman et al., 2017). In the model conditioning with paired triplets of stimuli shows a more potent conditioning effect, which became evident for a range of lower intensities where single stimuli were insufficient (Fig. 10).

### Cycle-Triggered Stimulation

To simulate cycle-triggered conditioning (Zanos et al., 2018), episodes of oscillatory beta activity were generated by modulating the bias input to Column B (episodes of 6 oscillations at 20 Hz occurred four times during each 10 second simulation time block). The local field potentials from Column B were filtered with a 15-25 Hz band pass and when this filtered LFP exceeded a given level, the next zero crossing was either taken as a 0° phase (when rising through zero) or 180° phase (falling through zero) to define the phase of the cycle trigger. During conditioning a stimulus was applied on all units in Column A whenever the cycle trigger for the specified phase occurred in B (Fig. 7A). To measure conditioning effects, we applied test pulses on Column A before and after the oscillatory episodes, as in the original experiments. These test pulses could cause a certain amount of tetanic conditioning, but this was lower than the changes caused by the cycle-triggered stimulation.

**Figure 7.**
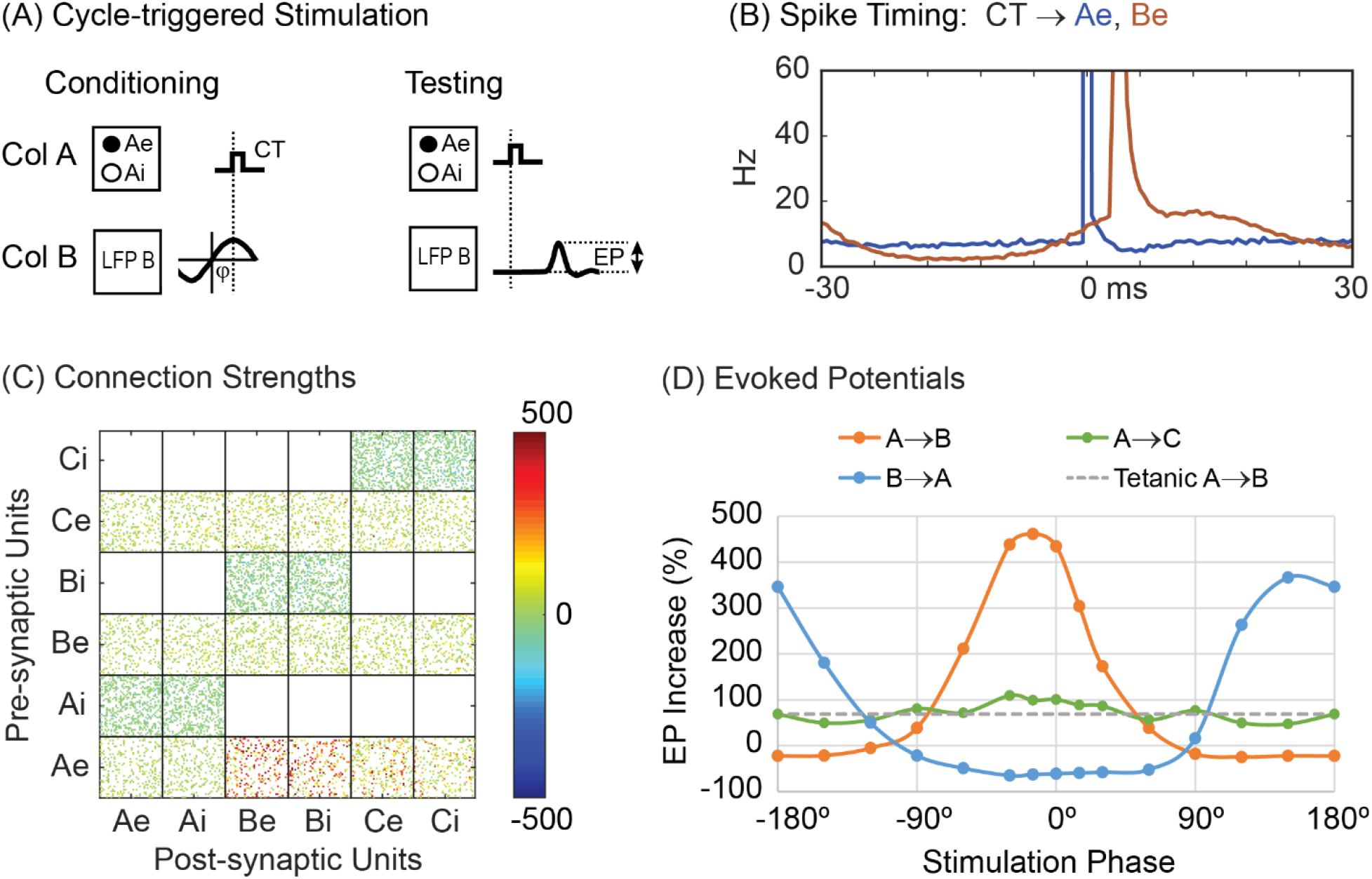
Cycle-Triggered Stimulation. (A) A stimulus is applied to Column A at a particular phase of local field potential oscillations in Column B. (B) Peristimulus time histogram of Ae and Be units during cycle-triggered stimulation at 0° phase. (C) Connection strengths after conditioning (0° phase). (D) Increase of the evoked responses as a function of stimulation phase. Legend identifies traces for stimulated and recorded columns. Dashed line shows A→B EP increase for tetanic stimulation on Column A with a rate approximately matching that for cycle-triggered stimulation.

As shown in Figure 7C, cycle-triggered stimulation at 0° phase shift increased connections from Column A→B and to a lesser extent the connections from Column A→C. In addition, the B→A connections were reduced. Cycle-triggered stimulation at 180° phase shift decreased connections from Column A→B and increased connections from Column B→A (Fig. 7D).

Figure 7B shows histograms of unit firings relative to the trigger for stimuli delivered at 0° phase shift. The Ae units fired in response to the stimuli (peak at zero in blue curve). The Be units (red) show the broad oscillatory increase and response to the stimuli.

Conditioning was also performed with stimuli delivered at other phases. Figure 7D shows how evoked responses vary with stimulation phase for A→B, A→C, and B→A connections. Maximum evoked responses for A→B occurred at about −15°, which matches the phase difference between the bias input modulation and the resulting modulation in the filtered LFP of Column B. Maximum B→A EP responses occurred at about 150° phase shift. The A→C connections showed a very weak modulation, which is consistent with experimental results for control sites whose LFPs were not entrained with B.

### Control for tetanic stimulation

An important control for conditioning effects is the effect of a comparable amount of open-loop stimulation not triggered by preceding activity. To reduce the effects of such tetanic stimulation, the amount of external bias drive and the area difference between the weakening and strengthening sides of the STDP curve were chosen such that tetanic stimulation, at rates similar to rates for conditioning protocols, did not cause large conditioning effects. The results of tetanic stimulation are shown in Figure 8, for Poisson-distributed stimulation delivered to Column B during the conditioning period. The connection matrices for tetanic stimulation show slight increases over the “No Stimulation” network in the connectivity from Column B to A and B to C (Fig. 8A). The changes in connectivity between columns as measured by EP amplitudes are illustrated in Figure 8B and plotted in Figure 8C as a function of stimulus frequency. Tetanic stimulation of B has the greatest effect on connections from B for frequencies between 8 and 16 Hz. The EP increases for other connections are small or slightly negative (Fig. 8B,C). Most of the conditioning protocols modified the A→B connections with triggered stimulation of Column B. In these cases the conditioned A→B effects were clearly larger than the tetanic effect which had the opposite effect of decreasing A→B EPs. For cycle-triggered stimulation of Column A, the conditioned effect was larger than the effect of tetanic stimulation of A alone (Fig. 7D).

**Figure 8.**
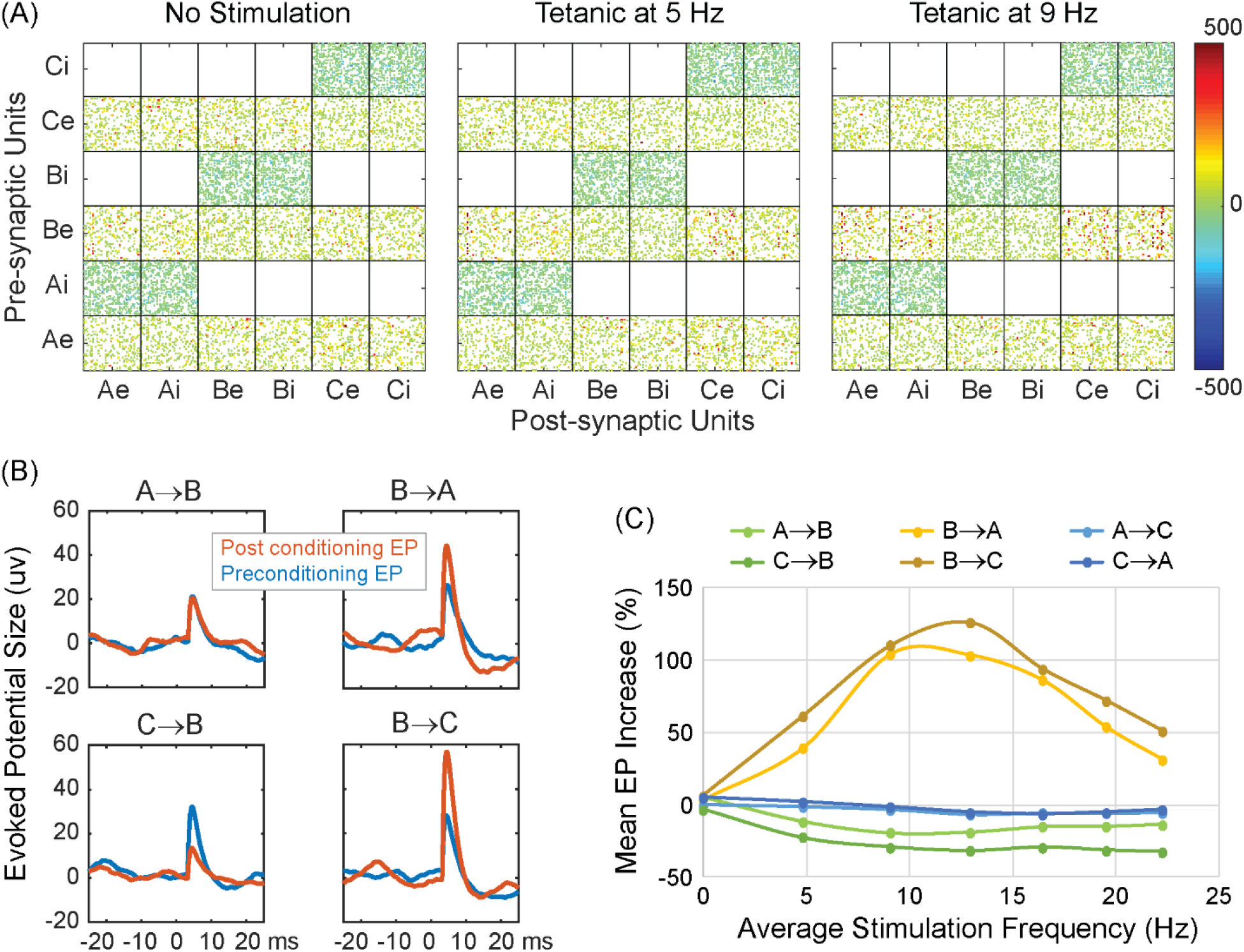
Tetanic Stimulation. (A) Effect on connection strengths of tetanic stimulation of Column B at 5 and 9 Hz (intensity of 2.0 mV), compared to no stimulation. (B) Examples of averaged EPs before and after tetanic stimulation of B at 9 Hz. (C) Mean (over 10 randomized networks) of averaged EP changes produced by tetanic stimulation of B at different frequencies. Legend identifies traces for test-stimulated → EP-recorded columns.

### Testable outcomes

Importantly, our IF model can simulate variables and procedures that were not documented in the original physiological experiments. Notably, the strength of synaptic connections in the model was documented by computing stimulus-evoked potentials (Figs. 4D, 5D, 6D, 7D, 8C), a test that could in principle be performed experimentally, but is technically difficult. Another comprehensive measure of conditioning effects on synaptic strength is the sum of all the synaptic weights connecting one column to another. There is no feasible physiological or anatomical experiment to measure these weights directly *in vivo*, but they can be readily calculated from the model. Significantly, Figure 9A shows that the computed weight changes closely track the changes in the EPs. Moreover, the weights can be tracked with high temporal resolution, during and after the conditioning to document the induction and decay of the conditioned weight changes. Figure 9B shows the time course of the weight changes in the A→B connections during and after a period of spike-triggered stimulation with STDP. The stimulation is applied during the first 4 seconds of each 10-second time block. The plot of the average time course over 50 blocks shows how the weights tend to increase during the stimulation and then decay towards their preconditioned state after stimulation is turned off. Figure 9C tracks these weight changes during and after two spike-triggered stimulation periods to show that weight changes can be induced and decay in a minute or two of simulated time. This figure also shows the variability in conditioning due to the training factor and firing rates of the units. A smaller training factor would reduce the variability of the conditioned weights but would also extend both the training and decay times.

**Figure 9.**
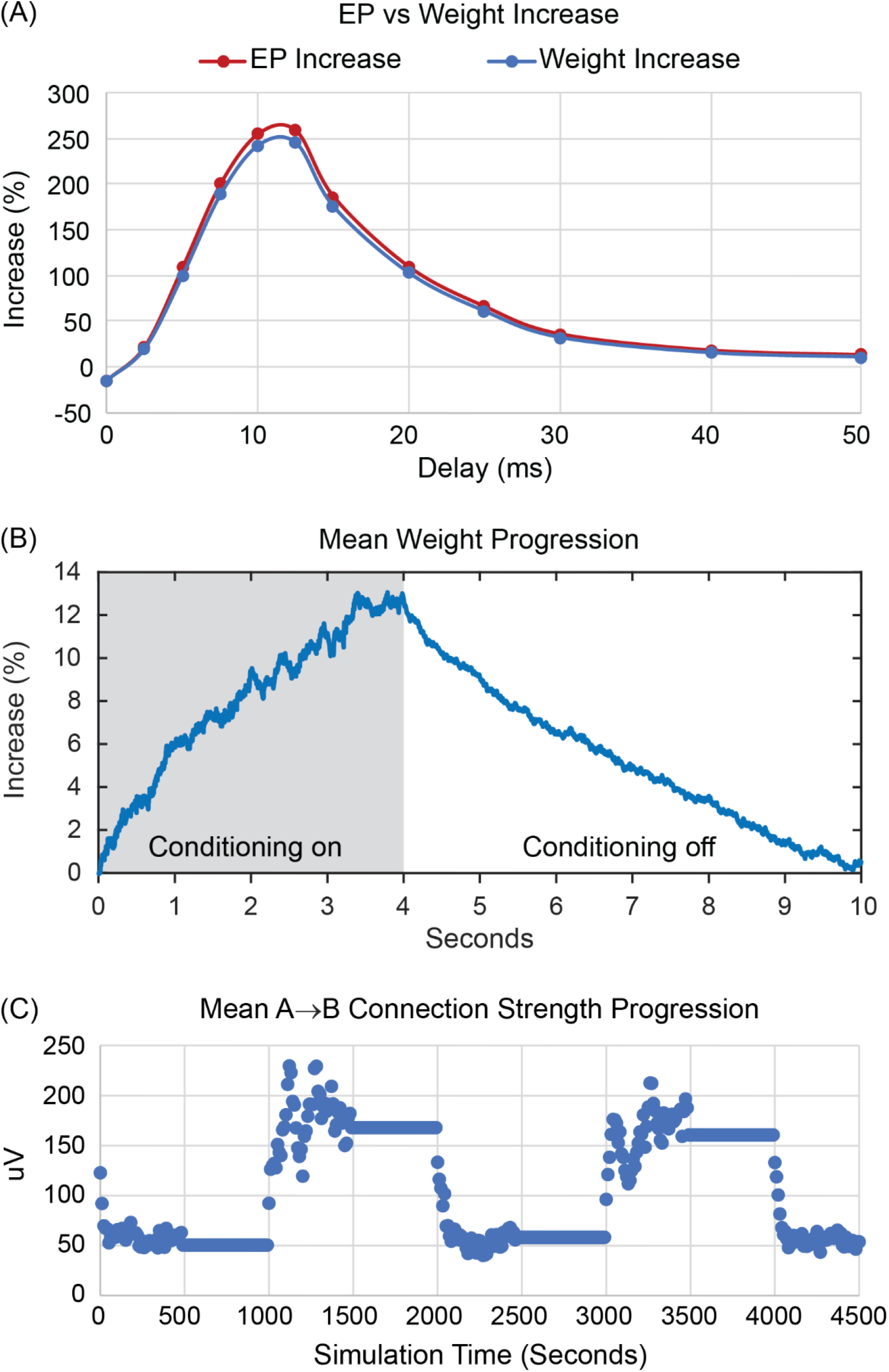
Network weight measurements. (A) Comparison between EP increase and mean A→B weight increase for conditioning with spike-triggered stimulation as function of spike-stimulus delay. (B) Average evolution of A→B weights during and after 4 sec of spike-triggered stimulation at 10 ms conditioning delay (averaged over 50 time blocks). (C) Mean A→B connection strengths calculated at the end of each 10 second time block and plotted over two passes through the pre-conditioning, testing, and spike-triggered stimulation periods.

Another testable property that the model can predict is the relative efficacy of conditioning as a function of stimulus intensity and number, as shown for the spike-triggered and paired pulse protocols in Figure 10. The curve for spike-triggered stimulus pairs rises faster than the curve for spike-triggered single stimuli; this is partly due to the pairs acting as a higher intensity stimulus, but that does not entirely explain the higher asymptote achieved. The curve for paired single-pulse stimulation initially rises more slowly than that for spike-triggered single stimuli, but then surpasses it at 2 mV, our standard conditioning simulation intensity. The curve for paired triplet stimuli (which had to be used experimentally) rises much faster than that for paired single-pulse stimulation, revealing a stimulus range over which the triplets are more effective than single pulses. The curves asymptote at high stimulus intensities when most of the units in the stimulated column respond.

**Figure 10.**
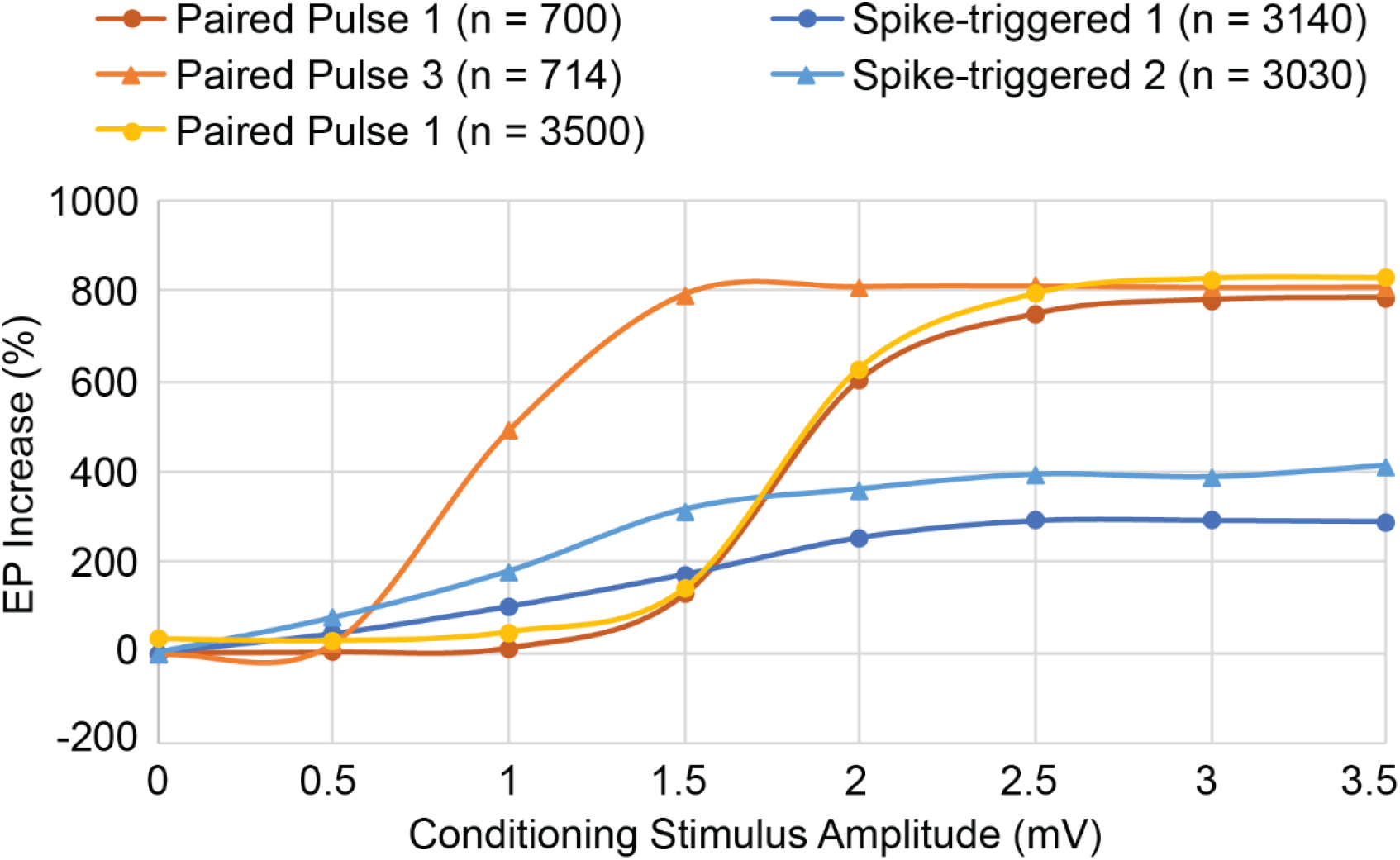
Effect of conditioning stimulus amplitudes and trains. Stimulating with pulse trains (33 ms interstimulus interval) instead of single pulses increases the efficacy of conditioning for spike-triggered stimulation (Spike-triggered 2 vs 1) and for paired pulse stimulation (Paired Pulse 3 vs 1) at all stimulus amplitudes. *n* is the total number of stimuli delivered during each simulation. Running the Paired Pulse 1 conditioning longer (n = 3500 compared to n = 700) does not provide the same increase in efficacy as Paired Pulse 3 (n = 714) until efficacy approaches an upper limit at stimulus amplitudes ≥ 2.5 mV.

### Testable predictions for novel protocols

Our IF model can also be used to predict outcomes of experiments not yet performed. For example, it is possible to trigger a pulse of *inhibition* of Column B from action potentials of a cell in Column A, which could be achieved by optogenetic spike-triggered inhibition (Fig. 11A). The inhibitory effect was modeled as a negative deflection (−2 mV) to the slow decaying part of the membrane potential. The results of this simulation show that the connections from A to B would be reduced (Fig. 11D). The trigger-aligned histograms in Figure 11B document the pairing of increases in Ae units with subsequent decreases in B units. The time-course of conditioned reduction in connections as a function of spike-pulse delay roughly resembles the inverse of the increase with spike-triggered stimulation.

**Figure 11.**
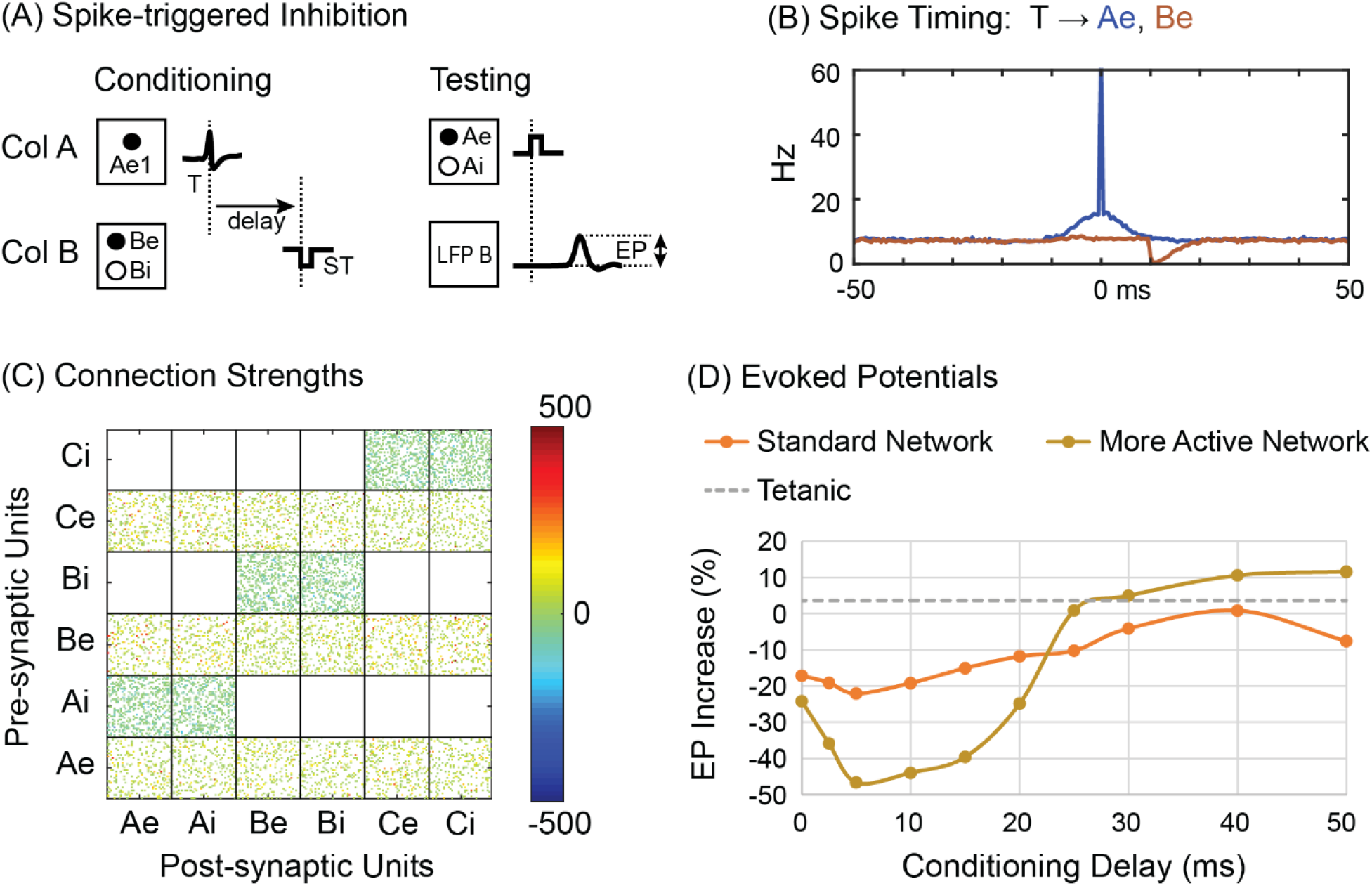
Spike-triggered inhibition. (A) Spikes in one unit in Column A (Ae1) trigger an inhibitory effect on all units in Column B. (B) Perispike histogram of Ae and Be units. (C) Post-conditioning connectivity matrix for 10 ms delay. (D) The standard network parameters yield a relatively mild EP decrease (orange curve). A more active network with 45% correlated bias drive, a weakening value of 0.5, and a −3mV stimulus produces a more robust post-stimulus inhibition (brown).

A second untested conditioning protocol involves triggering stimuli from threshold crossings of *gamma* range LFPs. On the rationale that gamma LFP reflects multiunit activity, these threshold crossings should detect coincident spiking activity that should also produce plasticity. Figure 12 shows the effects of stimuli in B triggered from band-passed mid-gamma LFP in Column A (in 50−80 Hz range, using 30% correlation in the bias spikes). The histogram of Ae unit spikes aligned with the falling gamma threshold crossings (Fig. 12B) shows a robust pre-stimulus peak in Ae units (and in LFP A) that generates the threshold crossing. The increase in A→B connections is shown in both the connection matrix (Fig. 12C) and in the size of EPs (Fig. 12D). Similar effects were found for triggers from high (80-100 Hz) and low range (30-50 Hz) gamma (Fig. 12D). Triggering from the rising slope of the gamma activity is associated with a later peak in Ae activity (Fig. 12B), consistent with a later latency and a smaller conditioning effect (Fig. 12D) than triggering from the falling slope.

**Figure 12.**
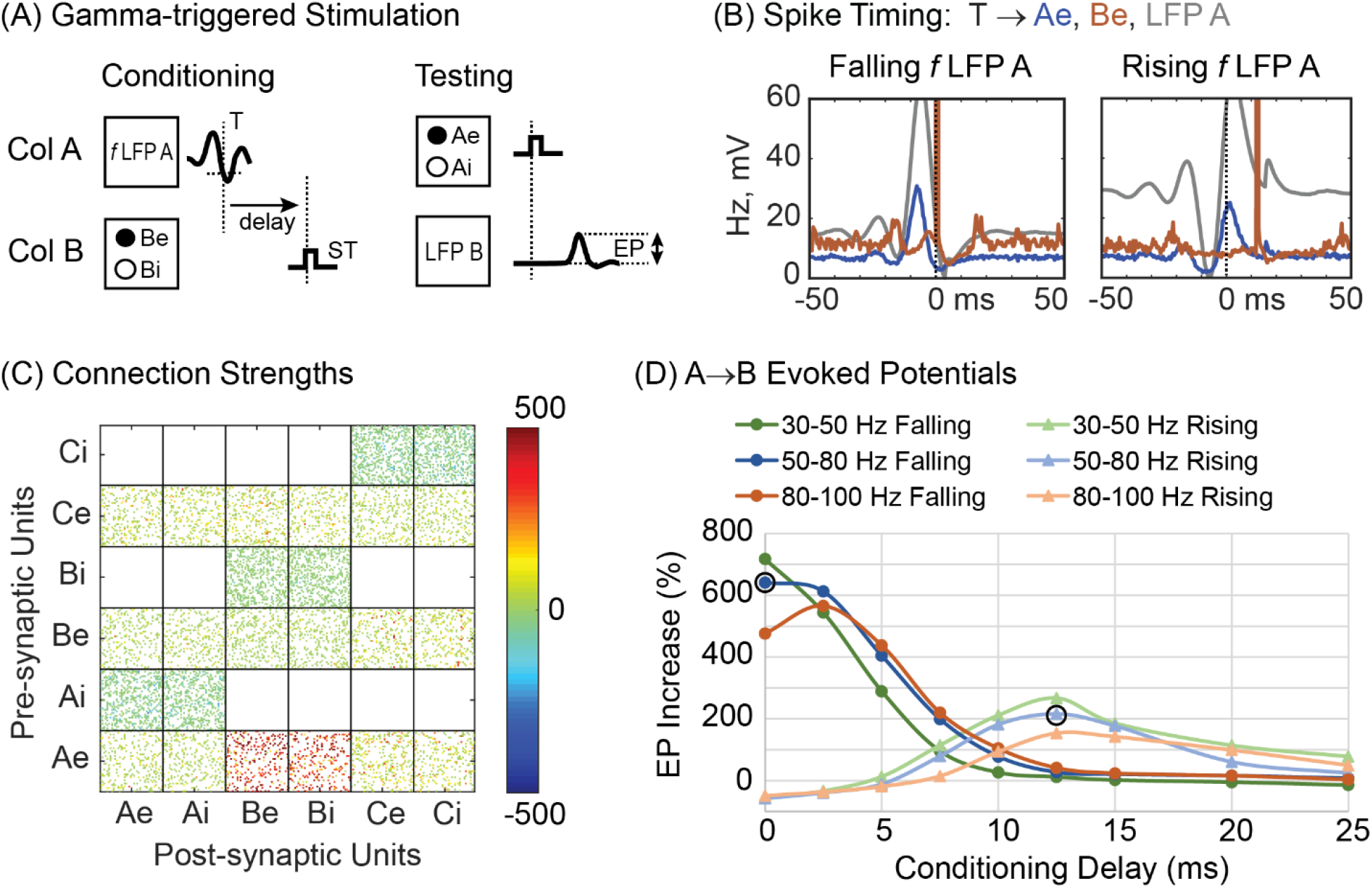
Gamma-triggered stimulation. (A) Stimuli were triggered from threshold crossings of LFP filtered in the gamma frequency ranges shown. (B) Firing rate histograms of Ae units (blue) and Be units (red) relative to triggers from falling and rising mid-range gamma filtered LFP in A. Grey curve shows simultaneous LFP in A. (C) Connection strength matrix for falling LFP filtered in mid gamma range (50-80 Hz). (D) changes in EPs for rising and falling threshold crossings for different gamma frequency ranges. Circled points used for histograms in (B).

### Relative efficacy of conditioning protocols

To compare the relative efficacy of different conditioning protocols, the resulting change of the A→B connections was tested by the change in the EP in B evoked by stimulating A. The results, plotted for different amplitudes of the conditioning stimuli, are shown in Figure 13. These simulations were run with the standard 30% correlated bias drive to each column. As a control for stimulation alone, the solid gray curve shows the effects of tetanic stimulation of Column B with exponentially distributed stimulation intervals, at the same average rate as the spike-triggered stimulation (blue curve). Above 1.5 mV the other four protocols all produce stronger effects than spike-triggered stimulation, probably because the stimuli involve larger numbers of Ae spikes. In particular, triggering from EMG and gamma produce larger effects than spike-triggered stimuli above 1 mV because they are selective for coincident Ae spikes (as shown in the histograms in Figure 5B and 12B).

**Figure 13.**
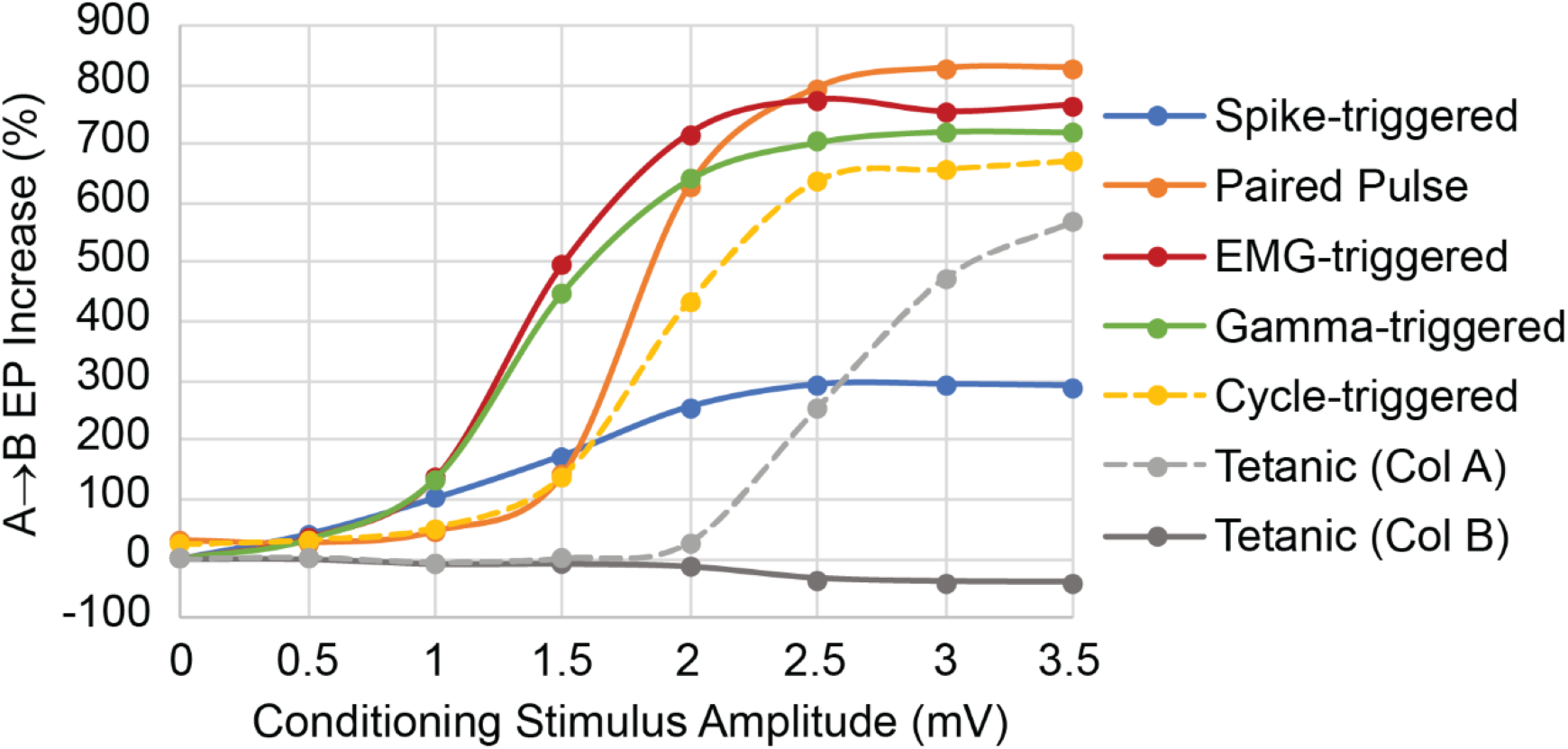
Relative efficacy of conditioning protocols. Curves show increase in A→B EP as function of stimulus intensity using standard network parameters. Tetanic stimulation was randomly spaced with rates approximating that for spike-triggered stimulation. Paired pulse interval was 10 ms, same as the spike-trigger delay, while rising EMG and falling gamma-trigger delay was 0 ms.

Tetanic stimulation of Column B (grey solid curve) is the control for spike-, gamma- and EMG- triggered stimulation of B. Tetanic stimulation of Column A (grey dashed curve) is the control for cycle-triggered stimulation of A (here performed at a phase of 0° in beta cycles). Both tetanic curves are relevant to paired pulse stimulation.

The conditioning effects asymptote at stimulus intensities above 2.5 mV, due to maximal activation of units in the stimulated column. The asymptote for spike-triggered stimulation (blue curve) is lower than those of the other conditioning protocols, but this was a function of the synchrony input driving the Ae units and could be increased by increasing the amount of synchronous relative to asynchronous input drive.

### Strengthening disynaptic pathways

The preceding conditioning protocols focused on changes induced in monosynaptic connections. The degree to which polysynaptic pathways can be modified would be of basic interest and clinical relevance for inducing targeted plasticity to treat lesions. Disynaptic conditioning was investigated for *paired stimulation* of A and B in the absence of direct connections between A and B (Fig. 14). The conditioning protocol is identical to that shown in Figure 6A for paired stimulation of intact networks, but now the most direct pathway between A and B is disynaptic, via Column C (Fig. 14A). The weight matrix shows clear increases in A→C and C→B connections after conditioning (Fig. 14C). This increase in the disynaptic path was a function of the intensity of the two stimuli, as shown by the increases in A→B EP size plotted for different combinations of intensity in Figure 14B. To produce conditioned changes with pairs of single pulses the stimulation intensity of A had to be at least 2.5 mV, higher than the typical 2 mV used in other simulations. This was likely necessary so that some tetanic conditioning from A→C would occur (as in Figure 13 Tetanic Col A). However, tetanic stimulation of A alone did not produce a sizeable disynaptic EP (Fig. 14B, B stim = 0). The A→B EPs increase appreciably for conditioning with interstimulus delays above 8 ms (Fig. 14D), about twice the minimal delay of 4 ms for direct connections (Fig. 6D).

**Figure 14.**
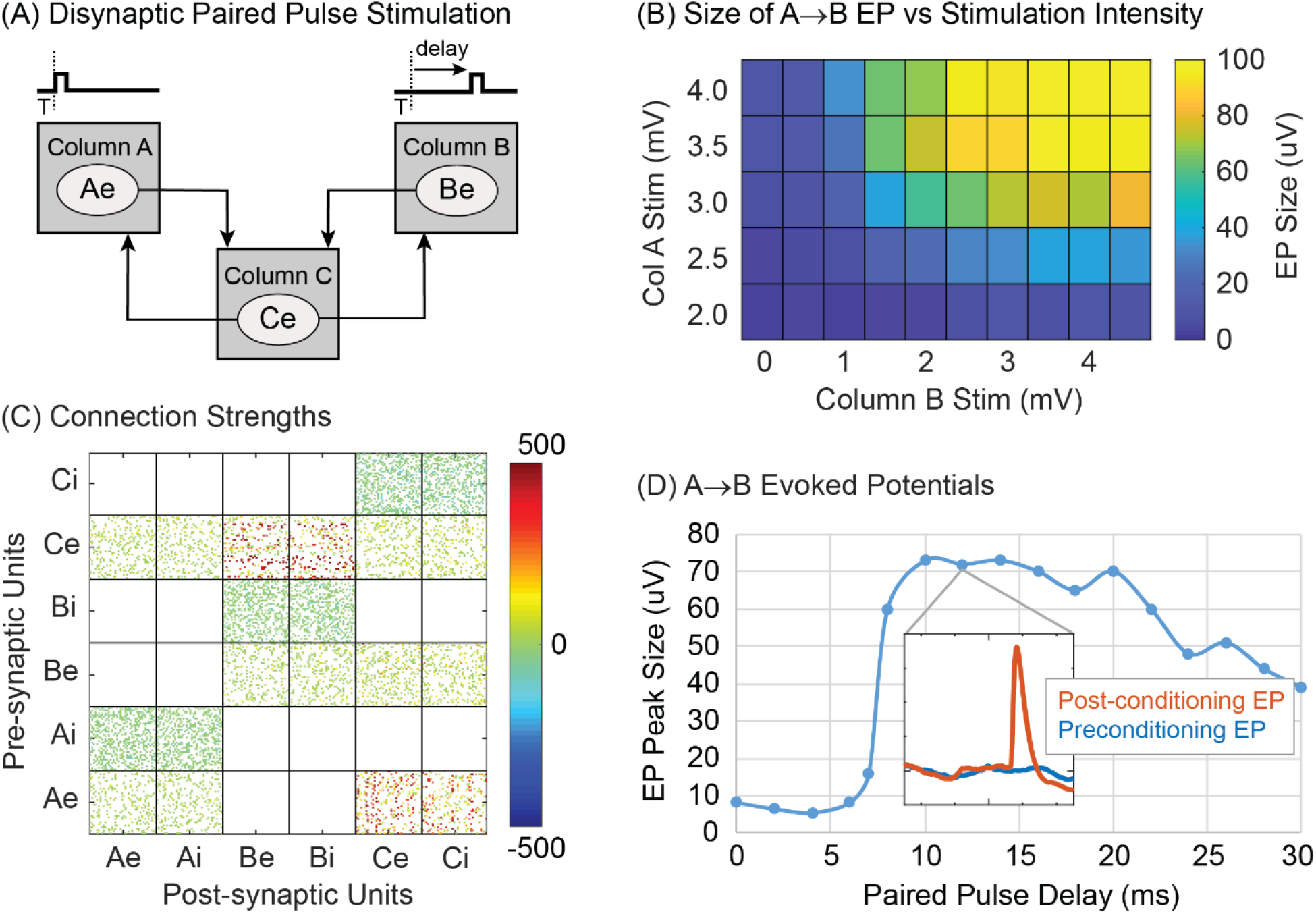
Disynaptic paired pulse stimulation. (A) Circuit connections between columns with standard A↔B connections removed. (B) Size of A→B EP as a function of stimulus intensities for the two pulses (for delay of 12 ms). (C) Post-conditioning connection matrix (for A and B stimulus intensities of 3 mV and delay of 12 ms). (D) Size of conditioned A→B EPs as a function of interstimulus delays (A and B stimulus intensities of 3 mV).

Although larger than standard stimuli were necessary to produce disynaptic conditioning with paired single pulses (Fig. 14B), paired triplet stimulation with standard intensities of 2 mV did produce clear disynaptic effects (Fig. 16).

*Cycle-triggered stimulation* could also be used to strengthen disynaptic connections (Fig. 15). The conditioning protocol is identical to normal cycle-triggered stimulation shown in Figure 7A, but now connections between Column A and Column B have been removed (Fig. 15A). Therefore, any EP in Column B caused by a stimulus in Column A must be mediated disynaptically, through Column C. Oscillatory drive was applied only to column B (Fig. 15B). The weight matrix again shows that conditioning produced clear increases in A→C and C→B connections (Fig. 15C). The stimulation intensity had to be increased to 3.5 mV, more than the typical 2.0 mV used in other simulations. This was probably necessary so that some tetanic conditioning from A→C would occur (as in Figure 13 Tetanic Col A). However, tetanic stimulation of A alone did not produce a sizeable disynaptic A→B EP (Fig. 15D, green line).

**Figure 15.**
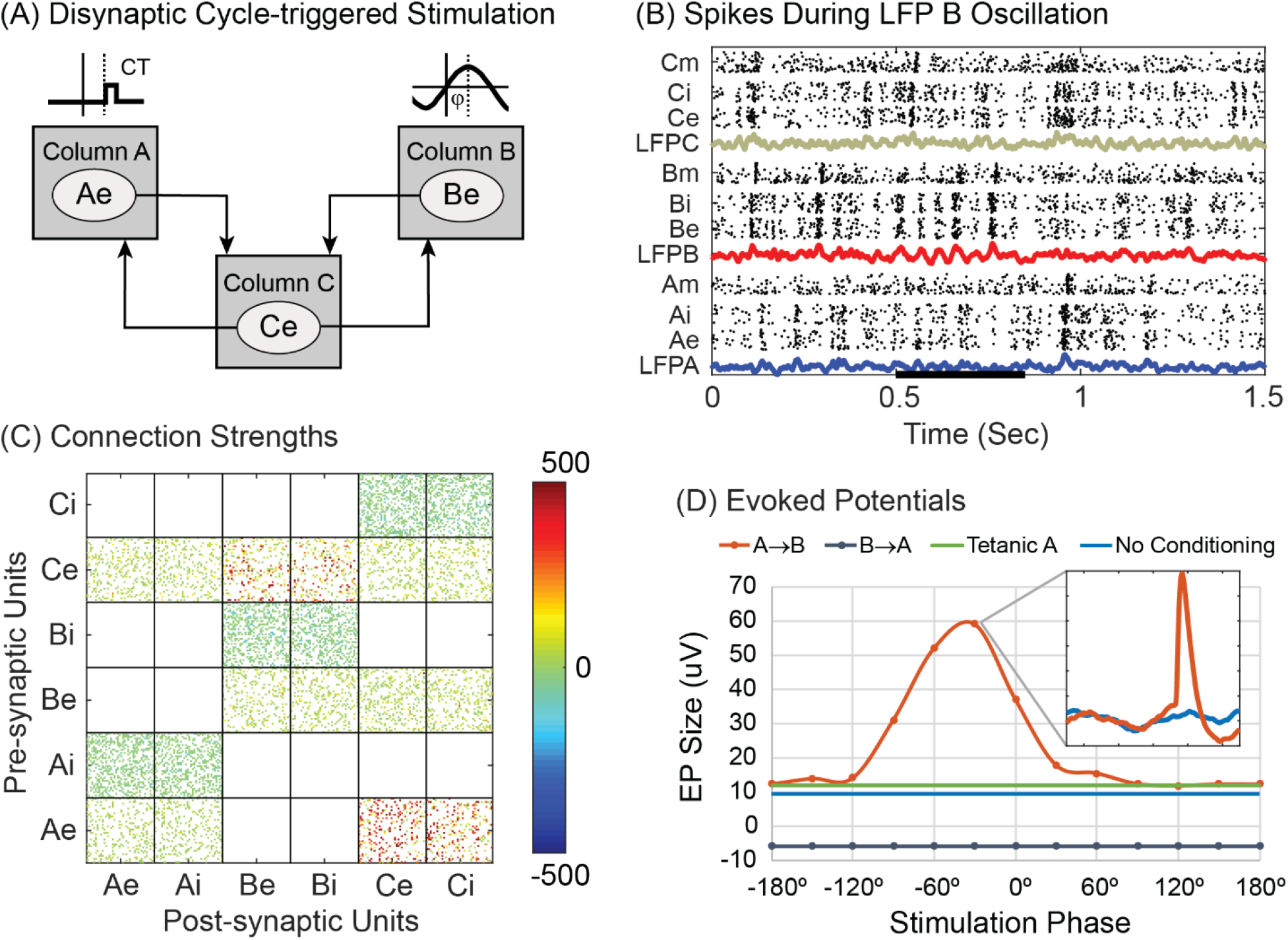
Disynaptic cycle-triggered stimulation. (A) Circuit connections between Columns A and B are removed. A stimulus (CT) on Column A is triggered on phases of an oscillatory episode in LFP B. (B) The oscillatory episode in Column B (starting at 0.5 sec) is produced by modulating the probability of external bias spikes to B with a sine wave (7 cycles at 20 Hz). (C) Post-conditioning connection matrix shows absence of A↔B connections, and the strengthened A→C and C→B connections that mediate the A→B EP. (D) Conditioned EP sizes for conditioning with different stimulation phases (φ). In contrast to cycle-triggered stimulation with full networks, the B→A EP was slightly depressed and unmodulated with phase. Inset shows A→B EP before (blue) and after (orange) conditioning.

**Figure 16.**
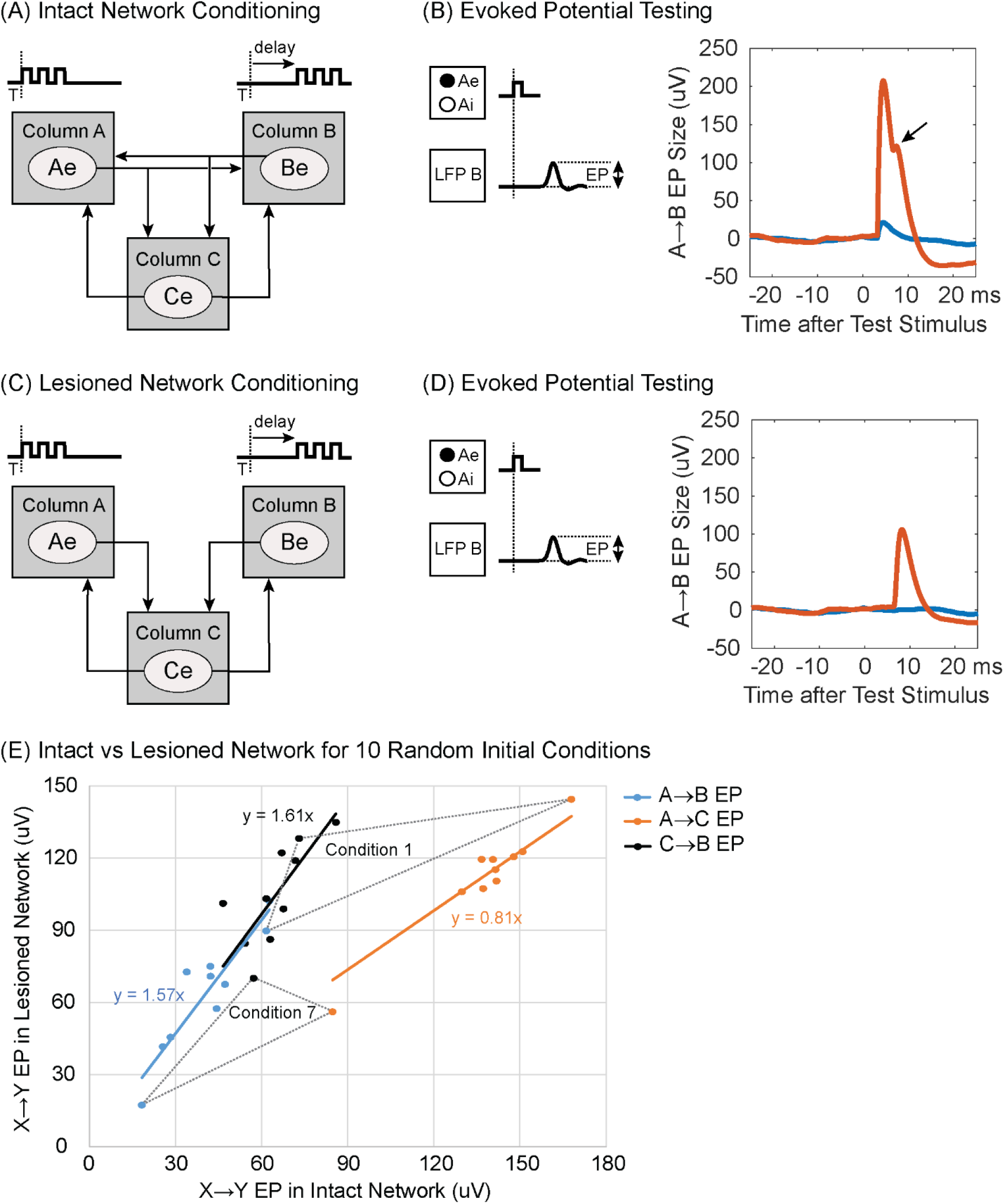
Paired triplet stimulation with standard 2 mV stimuli. (A) Intact network. (B) A→B EP after conditioning intact network with 10 ms delay between paired triplets (red) and before conditioning (blue). Arrow points to disynaptic component of EP. (C) Lesioned network trained without A↔B connections but other connections identical with intact network in (A). (D) A→B EP after conditioning lesioned network. (E) Sizes of intercolumn EPs in 10 conditioned intact and lesioned networks, with different starting weights. X→Y EP refer to monosynaptic A→C and C→B connections (orange and black points) and disynaptic A→C→B EPs (blue points); for the conditioned intact network the disynaptic A→C→B EPs were obtained after removing A↔B connections. Dotted triangles connect corresponding EPs from two starting conditions.

Disynaptic A→B connections could not be strengthened with spike- or gamma-triggered A→B stimulation, nor with stimulation of B triggered from EMG of Muscle A, even with larger than standard stimulus intensities. These protocols did strengthen A→C connections, but they did not produce the necessary increase in C→B connections.

The above tests involved disynaptic conditioning with A↔B connections deleted. A related question is whether comparable disynaptic conditioning occurs in intact networks as well. Intact networks conditioned with paired triplets using standard 2 mv stimuli (Fig. 16A) showed a clear disynaptic bump in the A→B EP (Fig. 16B, arrow). Lesioned networks without A↔B connections that were conditioned with the same paired triplets (Fig. 16C) showed a disynaptic A→C→B EP (Fig. 16D) whose timing coincided with the delayed bump produced in the intact network. However, their amplitudes differed. To measure the size of the disynaptic EP produced in the intact network we computed the A→C→B EP evoked with the A↔B connections removed for testing. The disynaptic EP produced by conditioning the lesioned networks was about 1.6 times larger than the recovered disynaptic component of the EP in the intact networks, with the identical conditioning protocol.

To investigate the reason for this difference we examined the sizes of disynaptic A→C→B EPs and the monosynaptic A→B and C→B EPs for the two networks. To elucidate the effects of initial weight choices on these values, this comparison was done for 10 intact networks, each starting with different initial synaptic weights and 10 lesioned networks that had identical initial weights except with A↔B connections deleted before conditioning (with paired triplets). Figure 16E plots the relative sizes of these EPs produced in the 10 networks. The disynaptic A→C→B EPs in the intact and lesioned networks (blue points) are proportional, with a linear slope of 1.57. The monosynaptic A→C and C→B connections (orange and black points) were also proportional, with slopes of 0.81 and 1.61, respectively. The two dotted triangles connect three corresponding EPs for two initial conditions (the ones that produced the largest and smallest EPs). The plots illustrate the variation in absolute sizes that can emerge from different starting points and also show that the sizes of the 3 EPs covary. These plots show that the larger disynaptic EP in the lesioned network can be attributed to a larger increase in the C→B connections.

## Discussion

Our IF model captures the experimental outcomes from four different conditioning protocols experimentally investigated in motor cortex of awake NHPs. A powerful feature of the model is to explore estimates of many variables that were not measured in the original experiments, such as changes in the connection matrices, sizes of evoked potentials (which measure the synaptic strength of inter-column connections), and peristimulus histograms of neural activity. Furthermore, the model predicts the outcomes of novel experiments not yet performed.

In comparison, a previous neural network model used populations of Poisson firing units and STDP to analytically compute the population effects produced by spike-triggered stimulation (Lajoie et al., 2017). That model replicated the experimental results on net changes in connectivity and showed that the amount of conditioning was dependent on the correlations between units. That model predicted that conditioning efficacy would be greater when cross-correlation peaks were wider, as they typically are during sleep compared to waking. In contrast, our IF model is based on simulating the synaptic connections between excitatory and inhibitory spiking neurons that integrate synaptic inputs to firing threshold. Our IF network provides cellular resolution of conditioning effects and their dependence on network parameters and simulates many different conditioning protocols.

A similar approach was used to model conditioning with spike-triggered stimulation in somatosensory cortex of NHPs (Song et al., 2013). That study examined the effect of spike-triggered and random microstimulation on firing rates and mutual information in S1. A large network of biophysical units with STDP reproduced many of the global physiological findings. In contrast, our smaller network of much simpler IF units was sufficient to replicate relatively detailed spatiotemporal measures of conditioning effects.

Other modelling studies have combined IF networks with STDP to investigate related issues. Adding a STDP rule to large populations of conductance-based units (Izhikevich, 2003) led to the emergence of interconnected groups under steady-state conditions (Izhikevich et al., 2004). Our model had fewer units, with predetermined connections between groups, and reached an equilibrium steady state under STDP before conditioning procedures were introduced. Other large-scale cortical network models showed that realistic synaptic plasticity rules coupled with homeostatic mechanisms led to the formation of neuronal assemblies that reflect previously experienced stimuli (Litwin-Kumar and Doiron, 2014) or recall sequences (Klos et al., 2018).

Bono and Clopath investigated STDP and dendritic spike mechanisms in producing plasticity in biophysically realistic neural models (Bono and Clopath, 2017). Another study showed that STDP rules had to be augmented with mechanism for heterosynaptic competition to generate networks capable of producing sequences of neural activity (Fiete et al., 2010). On the behavioral level, a model of orbitofrontal networks of IF units with STDP learned the rules of goal-directed actions (Koene and Hasselmo, 2005). These studies investigated various functional issues, in contrast to our focus on mechanisms of synaptic plasticity.

### Differences between the IF model and physiological mechanisms

It seems remarkable that our simple voltage-based IF model could replicate the results of many different physiological conditioning experiments. This robust performance raises the question of where the model has limitations that would inform further refinement. There are several differences between the performance of our model and the original physiological conditioning experiments. First, in the cortical spike-triggered stimulation experiments, not all site pairs developed strengthened connections (Jackson et al., 2006; Rebesco et al., 2010). This may simply be due to a lack of sufficient synaptic connections between those sites to strengthen. However, this explanation would not apply to the original paired stimulation experiments (Seeman et al., 2017), in which many pairs of sites did seem to have sufficient synaptic connections to mediate EPs, but these did not show conditioned changes.

Second, with paired stimulation *in vivo* a minimum of three pulses was required to produce conditioned changes (Rebesco and Miller, 2010; Seeman et al., 2017). In contrast, our IF model exhibited reliable conditioning even with pairs of single pulses, although triplet stimulation was substantially more potent than pairs of single stimuli over a range of intensities. The need for triplet instead of single pulses *in vivo* and the lack of conditioning effects for certain cortical sites remain to be better understood before these physiological observations can inform a change in the model.

A third difference between the model and physiological mechanisms concerns the duration of the conditioned effects. In vivo, the duration of effects was related to the amount of conditioning. The briefest conditioning effect (2 seconds) was produced by triggering stimuli from several cycles of beta oscillations (Zanos et al, 2018) and the longest conditioning effect (3 weeks) occurred in the behavioral recovery produced by 13 weeks of EMG-triggered spinal stimulation (McPherson, Miller et al. 2015). Other protocols involved intermediate durations of conditioning, and these produced correspondingly intermediate durations of effects (Jackson, Mavoori et al. 2006, Lucas and Fetz 2013, Nishimura, Perlmutter et al. 2013, Seeman, Mogen et al. 2017). In contrast, the time course of the induction and decay of conditioning effects in the model were similar for the different protocols (Fig. 9) and was determined by parameters such as training factor r, shape of STDP function and connectivity.

It seems possible that more sophisticated models, such as a network of biophysically more realistic units would help address some of these differences between the model and physiology. For example, the conductance-based units of Izhikevich capture biophysical properties that can replicate the complex firing properties of many types of cortical neurons in response to prolonged current injection (Izhikevich, 2003; Song et al., 2013). It will be interesting to use the conductance-based units with STDP to simulate the different conditioning protocols investigated here. This will require decisions about the number and connectivity of specific types of units in such a model as well as their biophysical parameters. The fact that our simple voltage-based IF network replicates the overall experimental observations may be related to two factors. It may represent an effective and sufficient average of the different types of biophysical neurons involved in the *in vivo* experiments. Second, the different response properties of biophysical units that can be demonstrated with prolonged intracellular or synaptic drive may be less critical for the phasic events that underly these conditioning protocols.

Our model used a STDP rule that required pre- and post-synaptic spikes to generate a change in synaptic connections. Physiological synapses could also be strengthened when the post-synaptic cell is merely depolarized after arrival of the presynaptic spike (Markram et al., 2012). This would represent a plasticity paradigm that does not require post-synaptic spikes. This mechanism would contribute to conditioning with cycle-triggered stimulation, for example. Intracellular recordings *in vivo* indicate that many cortical neurons show periodic membrane depolarization during beta oscillations that do not reach threshold for spiking (Chen D.F., 1993; Fetz et al., 2000). Phase-locked stimulation would modify the strength of synaptic connections to these depolarized neurons as well as to spiking neurons. Our simulations did not track the depolarization level of each unit, so the results of this STDP mechanism remain to be investigated in a future study.

There was a difference between the phase of maximum enhancement for A→B connections for cycle-triggered stimulation: the model showed a maximum at −15° (Fig. 7B) and experiments (Zanos et al, 2018) at around −90°. This difference could be due to differences in conduction times, stimulation and LFP dynamics, and physiological delays and plasticity mechanisms (above) not accounted for in the model.

### Parameter choices

The performance of our IF model depends of course on the choice of parameters. One choice involves the strength of baseline *synaptic connections*. To capture the small unitary synaptic potentials between cortical neurons (Matsumura et al., 1996; Markram et al., 1997) the plasticity parameters chosen for our model tend to produce a steady-state network with small connections when no activity-dependent conditioning protocol is in effect. This limits the effects produced by tetanic stimulation alone and makes conditioned increases in weights relatively robust. However, it also limits conditioned decreases in weights, since the starting size for most weights is already small. A possible future direction would be to develop a method to encourage a steady state of moderate weights so that both strengthening and weakening effects can manifest more equitably. This might also allow for more effects to emerge from inhibitory units, which play very little role in the network simulations shown here. We found that IF networks that were run without inhibitory units showed conditioning responses similar to networks that had many inhibitory connections.

A second choice in our model was using the same *STDP function* for inhibitory and excitatory units. The STDP function for inhibitory neurons has been found to vary, depending on the specific types of source and target neurons (Caporale and Dan, 2008; Kullmann and Lamsa, 2011; Vogels et al., 2011; Markram et al., 2012). We decided to use the same STDP function for both, which has empirical support (Haas et al., 2006), and to leave the consequences for different functions for inhibitory units, such as symmetric functions, to be explored in another study. Moreover, our STDP rule is based on pairs of pre- and post-synaptic spikes; models with “multi-spike” STDP interactions can produce different network dynamics (Babadi and Abbott, 2016).

A third choice in our model involved the relative amount of *external vs intracolumnar drive* on units. The input from local recurrent connections is a small fraction of a unit’s total input, which is dominated by external drive. Thus each “column” should be understood as a group of units with similar relations to the recording and stimulation parameters of the conditioning protocols rather than a network of units with sufficient self-connections to sustain substantial dynamics. Other studies have described the network dynamics emerging in populations of more strongly interconnected IF units (Brunel 2000, Izhikevich, Gally et al. 2004, Grytskyy, Tetzlaff et al. 2013, Wieland, Bernardi et al. 2015, Bos, Diesmann et al. 2016) but the focus of our simulations was on the effects of conditioning protocols on the involved units. Note that the online code for our model does allow exploration of different connection strategies.

A fourth choice in our model involved the relative amount of external *correlated input* bias. High amounts of correlated input can cause sufficient synchrony to strengthen connections from that column even in the absence of conditioning protocols. As the percent of correlated biases increases, the weakening portion of the STDP curve may be increased to prevent this effect from interfering with the manifestation of conditioning effects. For example, in our networks, the weakening value of 0.55 showed a spontaneous conditioning effect with 50% correlated/50% uncorrelated bias inputs, but did not show this effect with our chosen 30% correlated biases.

We found that networks run with many strong and fixed within-column weights could show conditioning effects with reduced or no correlated bias input. For example, by fully connecting within-column units with connections of 200 to 300 μV and preventing the STDP rule from modifying them, the various triggered stimulation protocols will cause conditioning without any external correlated input bias. These networks also show a function for inhibition to prevent run-away recurrent activity. However, such selective interventions to artificially boost the intracolumn connections introduce an inconsistency in the network mechanisms by artificially designating particular fixed connections and are therefore not included in this report. These simulations can be set up and explored with the online model.

In general, our choice of network parameters was guided by achieving realistic physiological performance. Notably, the same set of parameters was used to simulate all four conditioning protocols. Network performance was generally stable for modest deviations from the chosen parameters. The consequences of larger variations in different parameters would be interesting to explore but are beyond the scope of this report.

### Modeled connectivity and activity

An informative use of the model is to simulate activity and network connectivity that were not measured in the original experiments but that elucidate associated mechanisms. For example, calculating the EPs provides an easy and experimentally feasible measure of net connectivity between sites and can reveal its dependence on conditioning parameters like stimulus amplitude and delay, synchrony, etc. The changes to connection weights themselves can be computed in the model (but not experimentally); these weights closely tracked the EP measure (Fig. 9A). The weights further revealed a better time-resolved prediction of the induction and decay of these synaptic connections (Fig. 9B).

A particularly useful insight from the model is the activity of relevant units around the stimulus trigger events. Their stimulus-aligned histograms reveal why EMG-triggered and gamma-triggered stimulation are such effective conditioning protocols: they both select for coincident unit activity, as shown by the preceding peak in activity of A units (Figs. 5B, 12B). This peak preceding the stimulus-evoked post-synaptic activity in B is responsible for the strengthening of the A→B connections.

### Testable predictions

A possible therapeutic application of closed-loop conditioning is to strengthen polysynaptic pathways as well as strengthen direct connections. Such targeted plasticity could restore functional pathways lost to injury or stroke (Guggenmos et al., 2013). The model was used to examine conditioning of *disynaptic* links by performing A→B paired-pulse stimulation in a network without direct A→B connections and looking for development of A→C→B connections. This did occur, as shown by development of relevant disynaptic evoked potentials after conditioning and by the connectivity matrix (Fig. 14). Similarly, cycle-triggered stimulation could also strengthen disynaptic connections (Fig. 15); however, this protocol would be significantly more challenging to apply *in vivo* than paired stimulation. Both protocols involved sufficient stimulation of A to strengthen A→C connections. The other 3 protocols involved only stimulation of B and did not strengthen disynaptic A→C→B links. Another significant prediction of the model is that disynaptic effects are conditioned more robustly between sites that lack direct connections as compared to sites that also have direct connections. These simulations provide useful predictions of effective targeted plasticity paradigms and parameters to strengthen polysynaptic pathways.

The model also predicts several conditioning protocols that improve on methods used to date. Instead of spike-triggered single stimuli, the use of spike-triggered *bursts of stimuli* is more effective, increasing with the number of stimuli (Fig. 10). Second, *gamma-triggered stimulation* is more effective than spike-triggered stimulation because it detects coincident spikes that produce the gamma threshold crossing. Importantly, gamma-triggered stimulation is experimentally much easier to perform *in vivo* because it does not require long-term isolation of single action potentials. An important caveat here is that the model assumes that gamma is generated in a large number of A units that are synchronized and also connected to the B units. The efficacy of gamma-triggered stimulation would decrease if fewer units generating the gamma LFP in A sent connections to the stimulated units. Preliminary tests of this protocol by R. Yun (unpublished) indicate that conditioning effects are less robust *in vivo* in NHP motor cortex than in the model. This indicates that the connectivity of our network, designed to separate the recorded, stimulated and control groups may be too simple to accurately predict experimental outcomes for more complex biological networks in which these functional groups are intermingled with many neurons with other connectivities.

### Concluding comments

The simple IF spiking network described here has proven remarkably effective in capturing experimental results previously obtained in NHPs with four different conditioning protocols, including three with closed-loop activity-dependent stimulation. In addition to replicating the observed phenomena, the model also allows computation of underlying network behavior and correlated changes not originally documented. The model also makes significant predictions about protocols not yet investigated, including triggering bursts instead of single stimuli and gamma-triggered stimulation. The success of these simulations suggests that a simple voltage-based IF model incorporating STDP is sufficient to capture the essential mechanisms that produce targeted plasticity. Further detailed comparisons with physiological experiments will likely inform development of models with more realistic connectivity and biophysical properties.

